# Ancient *Clostridium* DNA and variants of tetanus neurotoxins associated with human archaeological remains

**DOI:** 10.1101/2022.06.30.498301

**Authors:** Harold P. Hodgins, Pengsheng Chen, Briallen Lobb, Xin Wei, Benjamin JM Tremblay, Michael J. Mansfield, Victoria CY Lee, Pyung-Gang Lee, Jeffrey Coffin, Ana T. Duggan, Alexis E. Dolphin, Gabriel Renaud, Min Dong, Andrew C. Doxey

## Abstract

The analysis of microbial genomes from human archaeological samples offers a historic snapshot of ancient pathogens and provides insights into the origins of modern infectious diseases. Here, through a large-scale metagenomic analysis of archeological samples, we discovered bacterial species related to modern-day *Clostridium tetani*, which produces the tetanus neurotoxin (TeNT) and causes the disease tetanus. We assembled draft genomes from 38 distinct human archeological samples spanning five continents and dating to as early as ~4000 BCE. These genomes had varying levels of completeness and a subset of them displayed hallmarks of ancient DNA damage. While 24 fall into known *C. tetani* clades, phylogenetic analysis revealed novel *C. tetani* lineages, as well as two novel *Clostridium* species (“*Clostridium* sp. X and Y”) closely related to *C. tetani*. Within these genomes, we found 13 TeNT variants with unique substitution profiles, including a subgroup of TeNT variants found exclusively in ancient samples from South America. We experimentally tested a TeNT variant selected from a ~6000-year-old Chilean mummy sample and found that it induced tetanus muscle paralysis in mice with potency comparable to modern TeNT. Our work identifies neurotoxigenic *C. tetani* in ancient DNA, new *Clostridium* species unique to ancient human samples, and a novel variant of TeNT that can cause disease in mammals.

## INTRODUCTION

*Clostridium tetani*, the causative agent of the neuroparalytic disease tetanus, is an important bacterial pathogen of humans and animals. After its spores enter wounds, it germinates, diffuses through oxygen-depleted and necrotic tissue, and produces a highly potent neurotoxin (tetanus neurotoxin, TeNT) that paralyzes hosts (Megighian et al., 2021; Popoff, 2020) leading to spastic paralysis. This may present as a local or a systemic effect, which can result in death due to paralysis of respiratory muscles and subsequent respiratory failure. As a wound-associated infectious disease, tetanus is estimated to have plagued *Homo sapiens* throughout history. Accounts of tetanus-like diagnostic features (“lockjaw”) date back as far as the ancient Greeks through written descriptions by Hippocrates (c. 380 BCE) (Miles, 2009; Pappas et al., 2008) and the ancient Egyptians as seen in reports from the Edwin Smith papyrus (c. 1600 BCE) (Sanchez & Burridge, 2007).

*C. tetani* was first isolated and cultivated in 1889 (Kitasato, 1889) and an early isolate of *C. tetani* (the 1920 Harvard E88 strain) is still widely used as a reference today. Genome sequencing of the E88 reference strain revealed that it possesses a single ~2.8 Mb chromosome and a ~74 kb plasmid (Bruggemann et al., 2003). This genomic organization is largely maintained among all known strains of *C. tetani*, with different strains varying in plasmid size (Brüggemann et al., 2015; Chapeton-Montes et al., 2019; Cohen et al., 2017). The plasmid is critical to pathogenicity as it contains the key virulence genes including *tent* which encodes the neurotoxin and *colT* which encodes a collagenase enzyme involved in tissue degradation. Based on comparative genomic analysis, modern *C. tetani* strains cluster into two phylogenetically distinct clades (Chapeton-Montes et al., 2019), which are closely related and exhibit low genetic variation with average nucleotide identities of 96-99%. Similarly, the *tent* gene is extremely conserved and exhibits 98% to 100% nucleotide identity across all strains. Modern *C. tetani* genomes therefore offer a limited perspective on the true diversity of *C. tetani* and its evolutionary history as a human pathogen.

The sequencing and analysis of ancient DNA (aDNA) from archeological samples provides unprecedented access to ancestral genomic information, and insights into the origins and evolution of modern species. In addition to human DNA, a significant proportion of genetic material preserved within ancient specimens is of a microbial origin (Bos et al., 2019; Drancourt & Raoult, 2005). Ancient microbial DNA, including that from ancient pathogens that once impacted humans, can be found within mummified remains, paleofeces, bones, teeth, and dental calculus (Warinner et al., 2015). Ancient DNA can be distinguished from modern DNA based on signatures of ancient DNA damage (Briggs et al., 2007; Dabney et al., 2013). Specifically, damaged ancient DNA is associated with an increase rate of deaminated cytosine residues that accumulate near the ends of DNA molecules. Due to the misincorporation of thymine instead of uracil by polymerases during amplification, this results in an apparent increased frequency of C→T transitions at the beginning and G→A transitions at the ends of sequence fragments (Briggs et al., 2007; Dabney et al., 2013). Damage levels exceeding 10% are considered to be indicative of genuinely ancient DNA (Sawyer et al., 2012). However, damage levels also depend on whether the DNA extract has been treated with uracil-DNA-glycosylase (UDG), which destroys the signal in the case of full UDG treatment, or reduces the signal at the ends of DNA molecules in the case of partial UDG treatment (Rohland et al., 2015). Once authenticated, reconstructed ancient microbial genomes can be compared with modern strains to investigate the genomic ancestry and adaptations underlying the emergence of historical epidemic strains. Groundbreaking aDNA studies on the evolutionary origins and emergence of major infectious diseases have been carried out in recent years including studies of *Mycobacterium tuberculosis* (Bos et al., 2014), the plague bacterium *Yersinia pestis* (Bos et al., 2011), *Mycobacterium leprae* (Schuenemann et al., 2013), *Helicobacter pylori* (Maixner et al., 2016), hepatitis B (Mühlemann et al., 2018), and variola virus (Duggan et al., 2016).

Here, using a large-scale metagenomic data mining of millions of sequencing datasets, we report the discovery of novel *C. tetani* related genomes including neurotoxin genes in ancient human DNA samples. Some strains and neurotoxins are phylogenetically distinct from modern forms, and some strains show strong hallmarks of ancient DNA damage indicative of an ancient origin. We further demonstrated that a neurotoxin variant from a ~6,000 year old Chinchorro mummy sample produces tetanus like paralysis in mice with a potency comparable to modern tetanus neurotoxins. Our findings uncover a widespread occurrence of *C. tetani* and related species associated with aDNA samples, expanding our understanding of the evolution and diversity of this important human pathogen.

## RESULTS

### Identification and assembly of *C. tetani* related genomes from aDNA samples

To explore the evolution and diversity of *C. tetani*, we performed a large-scale search of the entire NCBI Sequence Read Archive (SRA; 10,432,849 datasets from 291,458 studies totaling ~18 petabytes June 8, 2021) for datasets potentially containing *C. tetani* DNA signatures. Since typical homology-based search methods (e.g., BLAST) could not be applied at such a large scale, we used the recently developed Sequence Taxonomic Analysis Tool (STAT) (Katz et al., 2021) to search the SRA and identified 136 sequencing datasets possessing the highest total *C. tetani* DNA content [*k*-mer abundance >23,000 reads, k=32 base pair fragments mapping to the *C. tetani* genome] (Figure 1A, Table S1). Our search identified 28 previously sequenced *C. tetani* genomes (which serve as positive controls), as well as 108 uncharacterized sequencing runs (79 of human origin) with high predicted levels of *C. tetani* DNA content. Unexpectedly, 76 (96.2%) of these are aDNA datasets collected from human archeological specimens (Figure 1A), with the remaining three datasets being from modern human gut microbiome samples.

These 76 ancient DNA datasets are sequencing runs derived from 38 distinct archeological samples, which include tooth samples from aboriginal inhabitants of the Canary Islands from the 7^th^ to 11^th^ centuries CE (Rodríguez-Varela et al., 2017), tooth samples from the Sanganji Shell Mound of the Jomon in Japan (~1044 BCE) (Kanzawa-Kiriyama et al., 2017), Egyptian mummy remains from ~1879 BCE to 53 CE (Neukamm et al., 2020), and ancient Chilean Chinchorro mummy remains from ~3889 BCE (Raghavan et al., 2015) (Table S2). The 38 aDNA samples vary in terms of sample type (31 tooth, 6 bone and 1 chest extract), burial practices (27 regular inhumation and 11 mummies), sequencing method (26 shotgun datasets and 12 bait-capture approaches), and DNA treatment [6 uracil DNA glycosylase (UDG)-treated, 5 partial-UDG-treated and 27 untreated samples], all of which needs to be taken into account for interpretation of downstream analysis (Table S2).

Although these archeological samples are of human origin, STAT analysis of the 38 DNA samples predicted a predominantly microbial composition (Figure S1). *C. tetani*-related DNA was consistently abundant among predicted microbial communities, detected at 13.8% average relative abundance (Figure S1, Table S3). *M. tuberculosis* and *Y. pestis* were also detected (Figure S1) in several datasets associated with bait-capture sequencing of *M. tuberculosis* and *Y. pestis* from archaeological samples (Bos et al., 2016; Namouchi et al., 2018; Sabin et al., 2020).

To further explore the putative *C. tetani* in aDNA samples, we performed metagenome assembly using MEGAHIT for each individual sample and taxonomically classified assembled contigs using both Kaiju (Menzel et al., 2016) and BLAST (Altschul et al., 1997) to identify those mapping unambiguously to *C. tetani* and not other bacterial species (Table S4, S5). A majority (73%) of the alignments between assembled contigs and reference *C. tetani* genomes had percentage identities exceeding 99% (Figure 1B). Ninety percent of the alignments had percentage identities exceeding 90%, suggesting that a large fraction of assembled contigs are highly similar to regions of modern *C. tetani* genomes.

For each of the ancient DNA samples, we binned together all *C. tetani* like contigs to result in 38 putative, <u>a</u>ncient DNA associated <u>c</u>lostridial genome <u>bins</u> or “acBins”. We then performed QC analysis of each acBin using CheckM (Parks et al., 2015) to estimate genome completeness and contamination (Figure 1B, Table S6). Twenty-two acBins were more than 50% complete and 11 were more than 70% complete. Thirty-five acBins had low (<10%) contamination (Table S6). acBins with lower genome completeness were associated with bait-capture sequence methods, which makes sense as these methods aimed to enrich other organisms (Figure S2). We also examined the acBins for potential strain heterogeneity using two independent approaches: CheckM estimation (Table S6) as well as quantification of per-base heterogeneity from mapped reads (Table S2). Estimates of strain variation from both approaches correlated significantly (*r* = 0.47, *p* = 0.003) (Figure S3). Five strains (Sanganji-A2-Tooth, Chinchorro-Mummy-Bone, SLC-France-Tooth, Karolva-Tooth, Chincha-UC12-24-Tooth) were identified as possessing higher estimated levels of strain variation, but all were below 6% (CheckM) and 1.1% (average base heterogeneity).

### A subset of *C. tetani* genomes from archaeological samples are of ancient origin

Using the tools MapDamage2 (Jónsson et al., 2013) and pyDamage (Borry et al., 2021), we then examined the 38 acBins for elevated C→T misincorporation rates at the ends of molecules, a characteristic pattern of aDNA damage (Briggs et al., 2007; Dabney et al., 2013). Since these patterns are known to be affected by UDG treatment, we examined damage rates separately for full-UDG, partial-UDG, and untreated samples (Figure 1C). As expected, we observed the highest damage rates in the untreated samples, and the lowest damage rates in the full UDG-treated samples, indicating that the damage rates have been suppressed in some samples by UDG-treatment. The damage rates calculated by MapDamage and PyDamage were highly similar with a Pearson correlation of *r* = 0.99 (Table S2). Damage plots for all samples are shown in Figure S4 with additional data available in Tables S7 and S8.

Overall, seven acBins possessed a damage rate (5’ C→T misincorporation rate) exceeding 10%, which is indicative of aDNA (Sawyer et al., 2012) (top 5 shown in Figure 1D). In addition, all of the acBins except one (“Chincha-UC12-12-Tooth”) were verified by pyDamage as containing ancient contigs with *q*-values < 0.01 (Table S8). The highest damage rate (17.9%) occurred in the acBin from the “Augsburg-Tooth” sample, which is the third oldest sample in our dataset (~2253 BCE), despite this sample being partially UDG-treated (Figure 1D). As controls, evidence of ancient DNA damage was also observed in the corresponding human mitochondrial DNA (mtDNA) from the same ancient samples (Figure S4, Table S2), but not for modern *C. tetani* samples (Figure 1C). Additionally, no damage was detected in the three human gut derived *C. tetani* bins identified by our screen.

In general, we observed a significant correlation between damage rates of acBin DNA and corresponding human mtDNA from the same sample (*R*^2^ = 0.46, *p* = 3.4E-06, two-sided Pearson), although human mtDNA rates were generally higher in magnitude (Figure S5). Damage rates were higher for non-capture datasets as these generally received no UDG treatment (Figure S6A), and higher for samples associated with regular inhumations than those from mummies (Figure S6B). We also observed a significant correlation between acBin damage level and sample age, but only for mummy-derived samples (*R*^2^ = 0.50, *p* = 0.014) (Figure S6C). Together, these data suggest that a subset of the acBins display evidence of ancient DNA damage and are plausibly of an ancient origin.

**Figure 1.**
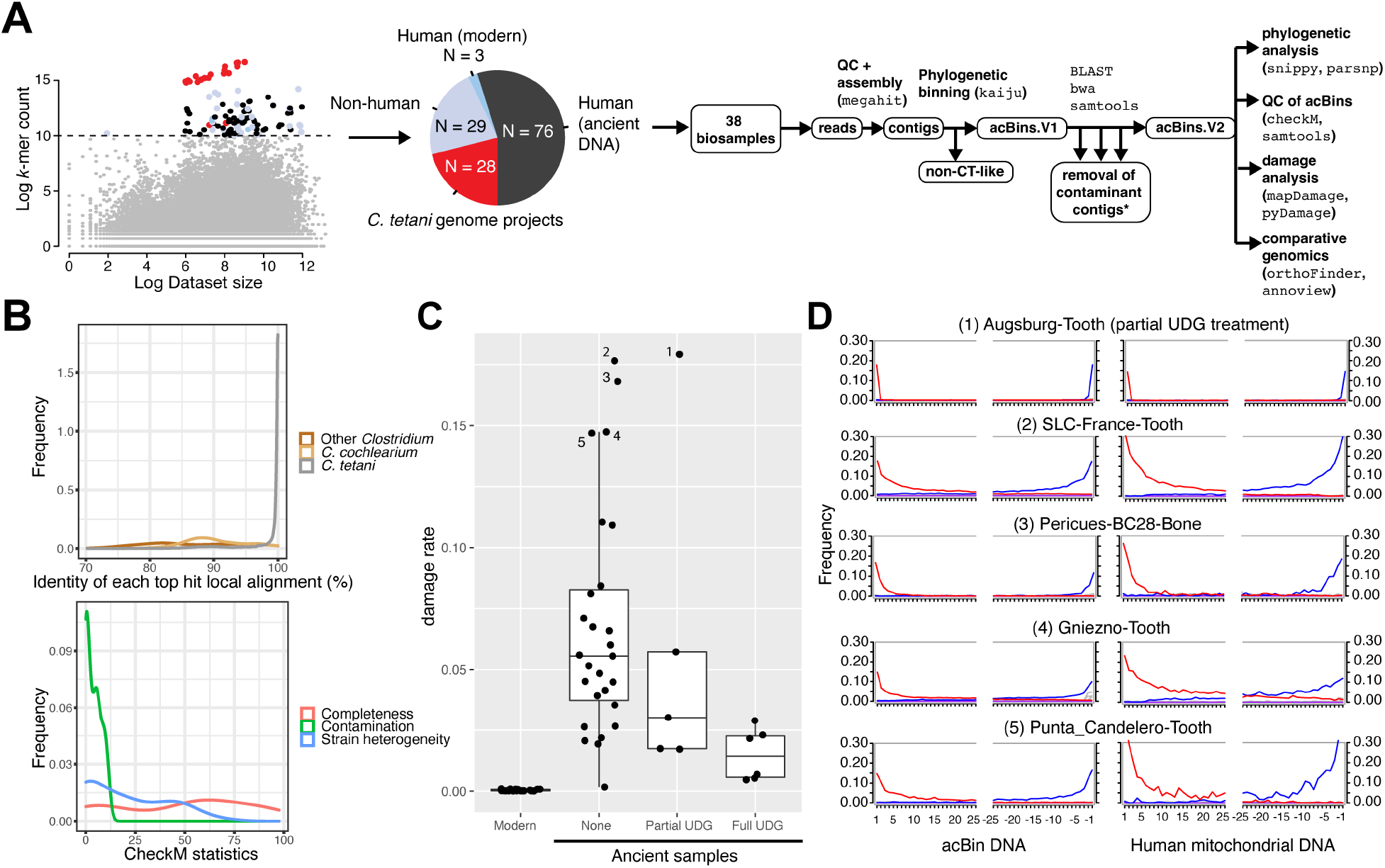
Petabase-scale screen of the NCBI sequence read archive reveals *C. tetani* related genomes in ancient human archeological samples. (A) General bioinformatic workflow starting with the analysis of 43,620 samples from the NCBI sequence read archive. Each sample is depicted according to its *C. tetani k*-mer abundance (y-axis) versus the natural log of the overall dataset size in megabases (x-axis). A threshold was used to distinguish samples with high detected *C. tetani* DNA content, and these data points are colored by sample origin: modern *C. tetani* genomes (red), non-human (light blue), modern human (blue), ancient human (black). The pie chart displays a breakdown of identified SRA samples with a high abundance of *C. tetani* DNA signatures. The 38 aDNA samples predicted to contain *C. tetani* DNA were further analyzed as shown in the bioinformatic pipeline on the right. (B) Top - density plot of the percentage identities of all BLAST local alignments detected between acBins and reference genomes including *C. tetani, C. cochlearium*, and other *Clostridium* sp. Bottom – density plot of the checkM results for the 38 acBins including estimated completeness, contamination, and strain heterogeneity levels. Completeness and contamination levels are percentage values. (C) MapDamage damage rates (5’ C→T misincorporation frequency) for 38 acBins subdivided by UDG treatment (none, partial, and full). Also shown are the damage rates for modern *C. tetani* genomes. (D) Damage plots for the top five acBins with the highest damage rates, and corresponding mtDNA damage plots. Shown is the frequency of C→T (red) and G→A (blue) misincorporations at the first and last 25 bases of sequence fragments. Increased misincorporation frequency at the edges of reads is characteristic of ancient DNA.

### Identification of novel *C. tetani* lineages and *Clostridium* species from ancient samples

To explore the phylogenetic relationships between the acBins and modern *C. tetani* strains, we first aligned their contigs to the reference *C. tetani* genome along with 41 existing, non-redundant *C. tetani* genomes (Chapeton-Montes et al., 2019), and clustered the genomes to produce a dendrogram (Figure 2A). Five acBins were omitted due to extremely low (<1%) genome coverage (see Methods), which could result in phylogenetic artifacts. We also included *C. cochlearium* as an outgroup, as it is the closest known related species to *C. tetani* based on phylogenomic analysis of available genomes (Mendler et al., 2019; Parks et al., 2018). Assessment of the genome-wide alignment for potential recombination showed no difference in estimated recombination levels for acBins compared to modern *C. tetani* genomes (Figure S7).

The genome-based dendrogram of the acBins and modern *C. tetani* strains (Figure 2A) matches the expected phylogenetic structure and contains all previously established *C. tetani* lineages (Chapeton-Montes et al., 2019). Ultimately, the acBins can be subdivided into those that cluster clearly within existing *C. tetani* lineages 1 or 2 and those that do not, which we have labeled “X” (8 acBins) and “Y” (1 acBin). Visualization of the acBin samples on the world map revealed a tendency for geographical clustering among acBins from the same phylogenetic lineage (Figure 2B). For example, lineage 1H acBins originate from ancient samples collected in the Americas, whereas most lineage 2 acBins originate outside of the Americas, and most clade X samples originate in Europe (Figure 2B).

#### acBins from C. tetani lineages 1 and 2

Twenty-four acBins fall within the *C. tetani* tree and possess average nucleotide identities (ANIs) of 96.4% to 99.7% to the E88 reference genome (Table S2), which is within the range considered to be the same species (Richter & Rosselló-Móra, 2009). These include new members of clades 1B (1 acBin), 1F (1 acBin), 1H (9 acBins), and 2 (9 acBins), expanding the known genomic diversity of clade 1H which previously contained a single strain and clade 2 which previously contained five strains (Figure 2A). Four additional acBins clustered generally within clade 1 but outside of established sublineages (Figure 2A).

Additionally, we used Parsnp (Treangen et al., 2014) to construct a more stringent, core SNP-based phylogeny from a reduced set of 11 acBins that could be aligned with high coverage to the reference *C. tetani* genome (see Methods) (Figure 2C, Figure S8). Only acBins from established *C. tetani* lineages 1 and 2 passed these criteria, and their phylogenetic positioning is consistent with their clustering pattern (Figure 2A). We also assembled a novel strain of *C. tetani* from a human gut sample (SRR10479805) which phylogenetically clustered with strain NCTC539 (98.7% average nucleotide identity; Table S9) from lineage 1G. The other two identified human gut samples were removed from further analysis as they predominantly matched *C. cochlearium* based on BLAST analysis.

#### acBins from clade “X” and branch “Y”

Nine acBins clustered outside of the *C. tetani* species clade. Eight of these cluster together as part of a divergent clade (labeled “X”) (Figure 2A). These samples span a large timeframe from ~2290 BCE to 1787 CE, are predominantly (7 of 8) of European origin (Figure 2B, Figure S9), and come from variable burial contexts including single cave burials, cemeteries, mass graves and burial pits (Andrades Valtueña et al., 2017; Bos et al., 2016; de la Fuente et al., 2018; Malmström et al., 2019; Stolarek et al., 2018; Susat et al., 2020; Valdiosera et al., 2018) (Table S2). Two of the samples from sites in Latvia and France are from plague (*Y. pestis*) victims (Bos et al., 2016; Susat et al., 2020), and another is from an individual with tuberculosis (Sabin et al., 2020). The highest quality clade X acBin is from sample “Augsburg-Tooth” (~2253 BCE), with 59.8% estimated completeness and 5.25% contamination (Table S6). Comparison of clade X acBins to other *Clostridium* species revealed that they are closer to *C. tetani* and *C. cochlearium* than any other *Clostridium* species available in the existing NCBI database, but are divergent enough to be considered a distinct species. On average, based on fastANI analysis of orthologous sequences (Jain et al., 2018) Clade X genomes have 86.3 +-1.8% ANI to *C. tetani* strain E88, and 85.2 +-1.6% ANI to *C. cochlearium* (Figure S10A, Table S10). Based on ANI analysis of the whole genome alignment, clade X genomes have 90.8 +-0.22% ANI to strain E88 (Table S2). These similarities were confirmed by analysis of BLAST alignment identities between clade X contigs and reference genomes (Figure S10B). As in the genome-wide tree, individual marker genes (*rpsL, rpsG* and *recA*) from clade X acBins also clustered as divergent branches distinct from *C. tetani* and *C. cochlearium* (Figure S11-S13). Finally, we re-examined the damage patterns according to phylogenetic clade, and found that clade X genomes possess the highest mean damage; 6/8 clade X genomes have a damage level exceeding 5% and 3/8 exceed 10% (Figure S6D, Table S2). These analyses suggest that clade X represents a previously unidentified species of *Clostridium*, including members of ancient origin. We designated this group *Clostridium* sp. X.

One sample (“GranCanaria-008-Tooth” from the Canary Islands dated to ~935 CE) also formed a single divergent branch (labeled “Y”) clustering outside all other *C. tetani* genomes (Figure 2A). Based on CheckM analysis, this acBin is of moderate quality with 74% completeness, and 0.47% contamination (Table S6). Comparison of the GranCanaria-008-Tooth acBin to the NCBI genome database revealed that it is closest to *C. tetani* and more distant to other available *Clostridium* genomes (Table S11). Based on fastANI (Jain et al., 2018), it exhibits an ANI of 87.4% to *C. tetani* E88, and 85.1% to *C. cochlearium*, below the 95% threshold typically used for species assignment (Table S11). Based on ANI analysis of the whole genome alignment, it has an 91.2% ANI to strain E88 (Table S2). To further investigate the phylogenetic position of this species, we built gene-based phylogenies with ribosomal marker genes *rpsL, rpsG* and *recA* (see Figures S11-S13). Each of these three genes support the GranCanaria-008-Tooth lineage as a divergent species distinct from *C. tetani*. The damage level for this acBin is relatively low (~4.0%), whereas its human mtDNA damage level is ~11.6% (Figure S4). We designated this acBin *Clostridium* sp. Y.

**Figure 2.**
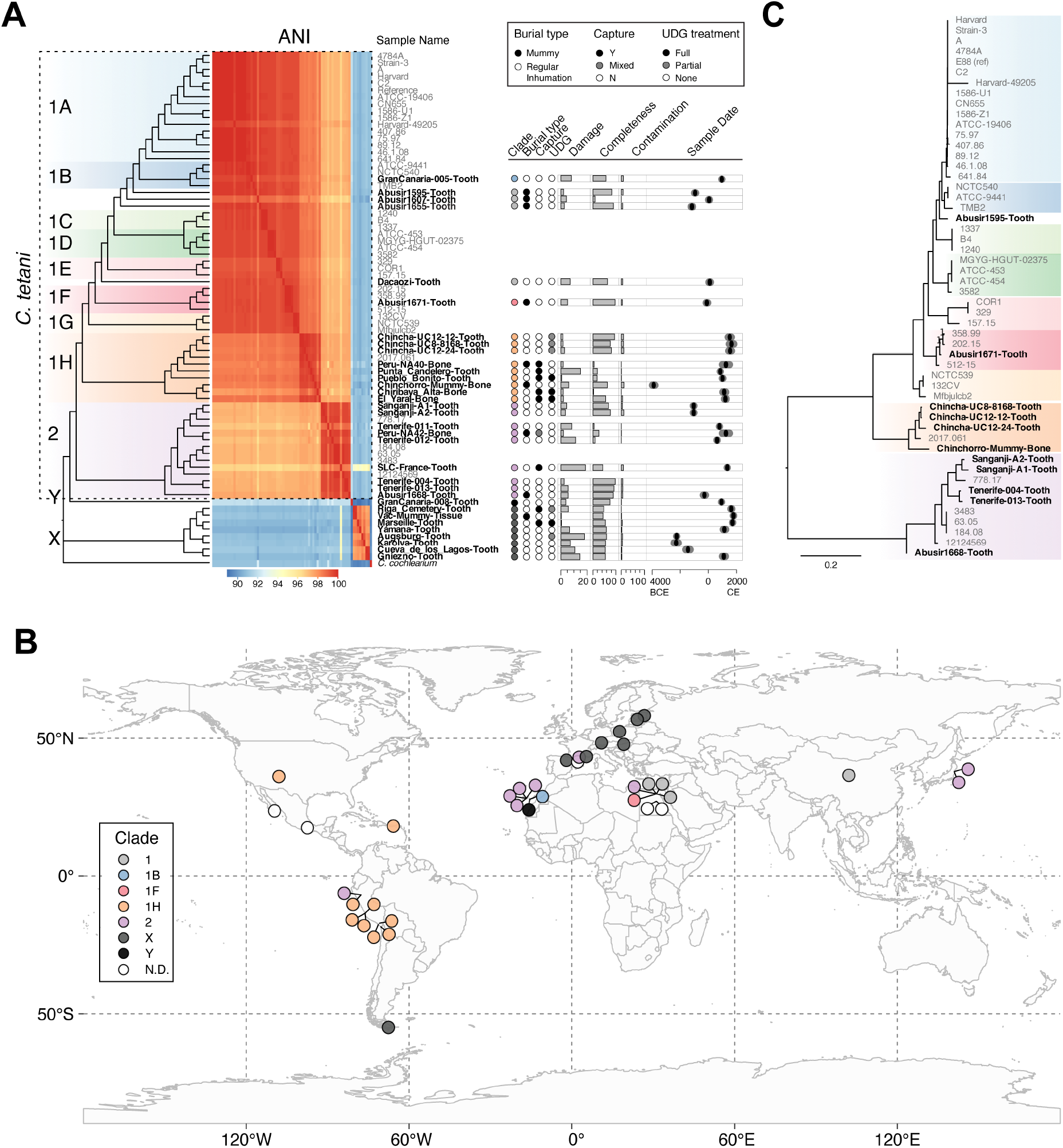
Phylogenetic analysis reveals known and novel lineages of *C. tetani* in ancient DNA samples, as well as a previously unidentified *Clostridium* species (“X”). (A) Dendrogram depicting relationships of acBins from ancient samples with modern *C. tetani* genomes. Novel branches are labeled “X” and “Y”, which are phylogenetically distinct from existing *C. tetani* genomes. Shown on the right of the dendrogram are metadata and statistics associated with each acBin including the estimated date of the associated archaeological sample. All metadata can be found in Table S2. (B) Geographic distribution of ancient DNA samples from which the 38 acBins were identified. Each sample is colored based on the acBin clustering pattern shown in A. (C) SNP-based phylogenetic tree of a subset of acBins from lineage 1 and 2 showing high similarity and coverage to the *C. tetani* reference genome. See Figure S8 for more details.

### Unique genomic differences discovered in *C. tetani* related genome bins from ancient samples

We next carried out a comprehensive comparison of genome content and structure between the acBins and modern *C. tetani* strains. We first clustered protein-coding sequences from all modern genomes and acBins into a set of 3,729 orthologous groups, and compared their presence/absence across all strains (see Methods). Based on this analysis, we observed considerable overlap in gene content between the acBins versus the modern reference genomes, with the greatest overlap observed between acBins from *C. tetani* lineages (1 and 2) and the smallest overlap observed for *Clostridium* sp. X (Figure S10C).

We then examined genome similarities by visualizing the alignment of each genome to the reference E88 chromosome and plasmid (Figure 3A). Several low coverage acBins can be seen in *C. tetani* lineages 1 and 2 (Figure 3A), which is expected given their low completeness estimates (Figure 2A). However, the divergent GranCanaria-008-Tooth genome (branch “Y”) and *Clostridium* sp. X consistently have a low alignment coverage, similar to that of *C. cochlearium* (Figure 3A), which we suspected may be due in part to these species being more distantly related to *C. tetani*. Consistent with the idea that clade X represents a distinct species from *C. tetani*, we identified fourteen genes that are unique to four or more clade X members and absent from all other *C. tetani* genomes. The genomic context of four of these genes (labeled by orthogroup) is shown in Figure S14. Although these genes are unique to clade X, their surrounding genes are conserved in other *C. tetani* genomes, implying that genome rearrangements may have resulted in these genes being either gained in *Clostridium* sp. X or lost in *C. tetani*.

To examine differences in plasmid gene content and structure directly, we then compared the gene neighborhoods surrounding the plasmid marker genes *repA* and *colT* (Figure 3C, expanded data shown in Figure S15). In several acBins from *C. tetani* lineages 1 or 2, the gene neighborhoods surrounding these genes are similar to that in modern strains (Figure S15). However, particularly in *Clostridium* sp. X and Y, we identified unique gene clusters distinct from those in modern strains. For example, in two *Clostridium* sp. X genomes and the *Clostridium* sp. Y genome, we identified a conserved toxin/antitoxin pair and a phage integrase flanking the *repA* gene (Figure 3C). We also observed a unique gene arrangement surrounding *colT* that is conserved in two clade X genomes (Figure S15). Additional differences were identified in a few lineage 2 acBins; for example, Tenerife-004-Tooth contains unique genes neighboring *repA*, and the Tenerife-013-Tooth acBin uniquely encodes the *repA* gene adjacent to its *tent* and *tetR* gene (Figure 3C).

We then performed a detailed comparison of the plasmid-encoded neurotoxin gene, *tent*, and its gene neighborhood (where possible) across the strains. As shown in Figure 3A as well as based on mapped read coverage to these regions (Figure 3B, Figure S16, Table S12), the *tent* gene was detected at relatively high depth of coverage in acBins from *C. tetani* lineages 1 and 2. The *tent* gene neighborhood structure from lineage 1 or 2 acBin strains is also similar or identical to that in modern strains, with the exception of Tenerife-013-Tooth (as it encodes the *repA* gene nearby) (Figure 3D)

However, in the acBins from lineage X and Y, the *tent* gene was either missing or was fragmented, suggesting a possible gene loss or pseudogenization event (Figure 3B). This pattern can be seen clearly in read coverage plots (Figure S16) and when normalizing *tent* depth of coverage to that of the plasmid-marker gene, *repA* (Figure 3B). The *tent* locus in the two *Clostridium* sp. X genomes for which assembly data is available over this region, appears to have undergone a deletion event resulting in the deletion of over 90% of the *tent* sequence as well as 3-4 neighboring genes (Figure 3D, Figure S17). This analysis further supports the idea that the *tent* fragment may be a non-functional pseudogene in these clade X strains.

Ultimately, our comparative genomic analysis of gene content and neighborhood structure demonstrates that the plasmids in several of our ancient samples (particularly those of *Clostridium* sp. X) are distinct from modern *C. tetani* plasmids, while the plasmids of acBins from lineages 1 and 2 are similar to those of existing *C. tetani* strains. This reinforces our earlier phylogenetic analysis indicating that clade X and branch Y represent a new *Clostridium* sp. that is closely related to but distinct from *C. tetani*.

**Figure 3.**
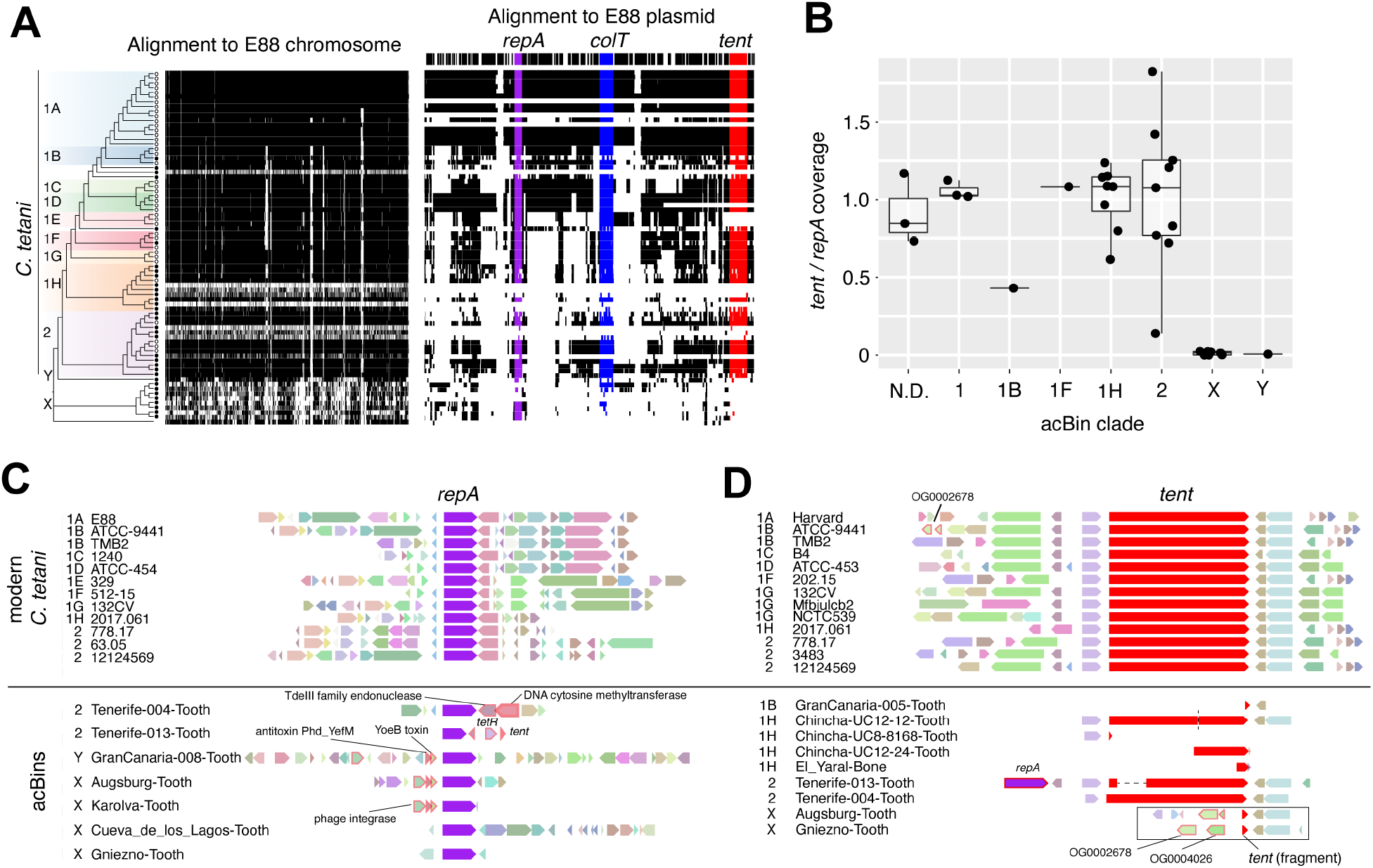
Comparative genomics of acBins versus modern *C. tetani* strains. (A) Visualization of the chromosomal and plasmid multiple sequence alignment. Orthologous blocks are shown in black and missing sequence is colored white. The reference gene locations are plotted above the alignments. (B) Per-clade coverage of the *tent* gene normalized to the coverage of *repA*. The coverage was calculated as the average depth of coverage based on mapped reads to each gene. See Figure S16 for the associated read pileups. (C) Gene neighborhoods surrounding *repA* genes in modern strains versus acBins. Selected unique differences identified in acBin gene neighborhoods are highlighted. (D) Gene neighborhoods surrounding *tent* genes in modern strains versus acBins. Selected unique differences identified in acBin gene neighborhoods are highlighted. The boxed region shows the assembled *tent* locus in two clade X acBins. Comparison reveals a putative deletion event in the clade X strains that has removed the majority of the *tent* gene along with five upstream genes, leaving behind conserved flanking regions. See Figure S17 for more information.

### Identification and experimental testing of novel TeNT variants

Given the considerable scientific and biomedical importance of clostridial neurotoxins, we next focused on *tent* and reconstructed a total of 18 *tent* gene sequences (all from lineage 1 and 2 acBins) from aDNA using a sensitive variant calling pipeline (see Methods). Six *tent* sequences have complete coverage, and 12 have 75-99.9% coverage (Table S13). Six partial *tent* sequences were also reconstructed but had lower average depth of coverage as shown in the read pileups (Figure S16). Four of the reconstructed *tent* sequences are identical to modern *tent* sequences, while 14 (including two identical sequences) are novel *tent* variants with 99.1-99.9% nucleotide identity to modern *tent*, comparable to the variation seen among modern *tent* genes (98.6-100%). We then built a phylogeny including the 18 *tent* genes from aDNA and all 12 modern *tent* sequences (Figure 4A). The *tent* genes clustered into three subgroups with modern and aDNA-associated *tent* genes found in subgroups 1 and 2, and aDNA-associated *tent* genes forming a novel subgroup 3 (Figure 4A). All three of the *tent* sequences in the novel *tent* subgroup 3 are from clade 1H aDNA strains.

We then visualized the uniqueness of aDNA-associated *tent* genes by mapping nucleotide substitutions onto the phylogeny (Figure 4B, Figure S18), and focusing on “unique” *tent* substitutions found only in ancient samples and not in modern *tent* sequences. We identified a total of 46 such substitutions that are completely unique to one or more aDNA-associated *tent* genes (Figure 4B, Figure S19, Table S14), which were statistically supported by the stringent variant calling pipeline (Table S15). The largest number of unique substitutions occurred in *tent*/Chinchorro from *tent* subgroup 3, which is the oldest sample in our dataset (“Chinchorro mummy bone”, ~3889 BCE). *tent*/Chinchorro possesses 18 unique substitutions not found in modern *tent*, and 12 of these are shared with *tent*/El-Yaral and 10 with *tent*/Chiribaya (Figure 4B). The three associated acBins also cluster as neighbors in the phylogenomic tree (Figure 2A), and the three associated archaeological samples originate from a similar geographic region in Peru and Chile (Figure S20). These shared patterns suggest a common evolutionary origin for these *C. tetani* strains and their unique neurotoxin genes and highlight *tent* subgroup 3 as a distinct group of *tent* variants exclusive to ancient samples (Figure 4A).

We then focused on *tent*/Chinchorro as a representative sequence of this group as its full-length gene sequence could be completely assembled. The 18 unique substitutions present in the *tent*/Chinchorro gene result in 12 unique amino acid substitutions, absent from modern TeNT protein sequences (L140S, E141K, P144T, S145N, A147T, T148P, T149I, P445T, P531Q, V653I, V806I, H924R) (Table S16). Seven of these substitutions are spatially clustered within a surface loop on the TeNT structure (Masuyer et al., 2017) and represent a potential mutation “hot spot” (Figure 4C). Interestingly, 7/12 amino acid substitutions found in TeNT/Chinchorro are also shared with TeNT/El-Yaral and 5/12 are shared with TeNT/Chiribaya (Table S16). As highlighted in Figure 4C, TeNT/Chinchorro and TeNT/El-Yaral share a divergent 9-aa segment (amino acids 141-149 in TeNT, P04958) that is distinct from all other TeNT sequences. Reads mapping to the *tent*/Chinchorro gene show a low damage level similar to that seen in the *C. tetani* contigs from this sample, and their damage pattern is weaker than the corresponding damage pattern from the associated human mitochondrial DNA (Figure 4D).

Given the phylogenetic novelty and unique pattern of substitutions observed for the *tent*/Chinchorro gene, we sought to determine whether it encodes an active tetanus neurotoxin. For biosafety reasons, we avoided the production of a *tent*/Chinchorro gene construct and instead used sortase-mediated ligation to produce limited quantities of full-length protein toxin (Figure S21), as done previously for other neurotoxins (Zhang et al., 2017, 2018). This involved producing two recombinant proteins in *E. coli*, one constituting the N-terminal fragment and another containing the C-terminal fragment of TeNT/Chinchorro, and then ligating these together using sortase. The resulting full-length TeNT/Chinchorro protein cleaved the canonical TeNT substrate, VAMP2, in cultured rat cortical neurons, and can be neutralized with anti-TeNT antisera (Figure 4E, Figure S21). TeNT/Chinchorro induced spastic paralysis *in vivo* in mice when injected to the hind leg muscle, which displayed a classic tetanus-like phenotype identical to that seen for wild-type TeNT (Figure 4F). Quantification of muscle rigidity following TeNT and TeNT/Chinchorro exposure demonstrated that TeNT/Chinchorro exhibits a potency that is indistinguishable from TeNT (Figure 4G). Together, these data demonstrate that the reconstructed *tent*/Chinchorro gene encodes an active and highly potent TeNT variant.

**Figure 4.**
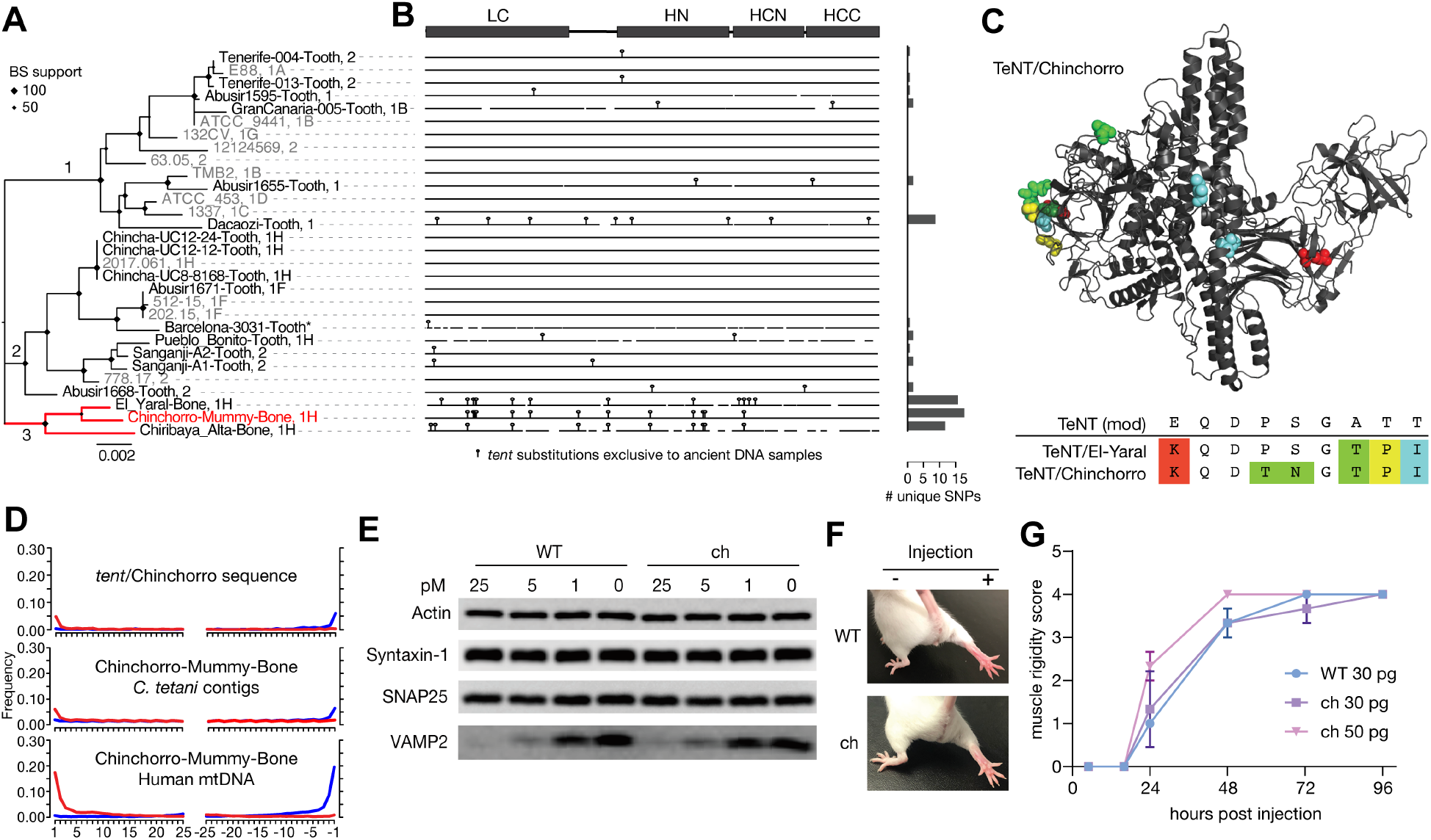
Analysis and experimental testing of a novel TeNT lineage identified from ancient DNA. (A) Maximum-likelihood phylogenetic tree of *tent* genes including novel *tent* sequences assembled from ancient DNA samples and a non-redundant set of *tent* sequences from existing strains in which duplicates have been removed (see Methods for details). The phylogeny has been subdivided into three subgroups. Sequences are labeled according to sample followed by their associated clade in the genome-based tree (Figure 2A), except for the Barcelona-3031-Tooth sequence (*) as it fell below the coverage threshold. (B) Visualization of *tent* sequence variation, with vertical bars representing nucleotide substitutions found uniquely in *tent* sequences from ancient DNA samples. On the right, a barplot is shown that indicates the number of unique substitutions found in each sequence, highlighting the uniqueness of subgroup 3. (C) Structural model of TeNT/Chinchorro indicating all of its unique amino acid substitutions, which are not observed in modern TeNT sequences. Also shown is a segment of the translated alignment for a specific N-terminal region of the TeNT protein (residues 141-149, Uniprot ID P04958). This sub-alignment illustrates a segment containing a high density of unique amino acid substitutions, four of which are shared in TeNT/El-Yaral and TeNT/Chinchorro. (D) MapDamage analysis of the *tent*/Chinchorro gene, and associated *C. tetani* contigs and mtDNA from the Chinchorro-Mummy-Bone sample. (E) Cultured rat cortical neurons were exposed to full-length toxins in culture medium at indicated concentration for 12 hrs. Cell lysates were analyzed by immunoblot. WT TeNT and TeNT/Chinchorro (“ch”) showed similar levels of activity in cleaving VAMP2 in neurons. (F-G) Full-length toxins ligated by sortase reaction were injected into the gastrocnemius muscles of the right hind limb of mice. Extent of muscle rigidity was monitored and scored for 4 days (means ± se; n=3 per group, 9 total). TeNT/Chinchorro (“ch”) induced typical spastic paralysis and showed a potency similar to WT TeNT.

## DISCUSSION

In this work, large-scale data mining of millions of existing genomic datasets revealed widespread occurrence of neurotoxigenic *C. tetani* and related species of *Clostridium* in aDNA samples from human archaeological remains. Our study provides three main findings: 1) the first identification of neurotoxigenic *C. tetani* from archaeological samples including several *C. tetani* strains of plausibly ancient origin; 2) the discovery of novel lineages of *C. tetani* as well as entirely new species of *Clostridium* (clade X and lineage Y); and 3) the identification of novel variants of TeNT including TeNT/Chinchorro which we demonstrate to be an active neurotoxin with extreme potency comparable to modern TeNT variants.

Our work is unique from previous studies of aDNA in several respects. First, using recent advances in petabase-scale genomic data-mining (Katz et al., 2021), we were able to perform a large-scale survey of sequencing datasets including all publicly available DNA samples in the NCBI SRA, greatly enhancing our ability to discover patterns across spatially and temporally diverse datasets. Because of the massive size of the NCBI SRA database, traditional sequence-alignment based methods such as BLAST are not computationally feasible. The STAT method (Katz et al., 2021) provided a heuristic approach to identify potential datasets from the SRA that could then be targeted for deeper analysis including metagenomic assembly and phylogenomics. Importantly, we did not specifically look for *C. tetani* in ancient DNA, but rather this came as an unexpected finding from the results of our large-scale screen. Also unexpected was the considerable diversity of ancient samples in which we identified neurotoxigenic *C. tetani*, which revealed a strong association between this organism (and related species) and human archaeological samples. Despite the abundance of environmental (e.g., soil metagenomic) samples in the SRA, these samples did not come to the surface of our genomic screen for *C. tetani*. This is consistent with the idea that, although *C. tetani* spores may be ubiquitous in terrestrial environments such as soil, these spores may be rare and so *C. tetani* DNA may not regularly appear at appreciable levels in shotgun metagenomes. Lastly, it is also noteworthy that the majority (31/38) of *C. tetani* aDNA samples in our study originated from teeth. Teeth are commonly used in aDNA studies due to the survival and concentration of endogenous aDNA content (Adler et al., 2011). The colonization of tooth and bone tissue by *C. tetani* is also not unprecedented, as infections of these tissues have been previously but infrequently reported in clinical cases (Darraj et al., 2017; Levy et al., 2014).

It is important to point out that the identification of neurotoxigenic *C. tetani* in aDNA samples alone is not sufficient to implicate tetanus as a cause of death or even suggest that the corresponding *C. tetani* strains are contemporaneous with the archaeological samples. A variety of environmental factors and mechanisms may account for the presence of toxigenic clostridia in aDNA samples, including the possibility of their introduction by human handling of archaeological samples after death, or post-mortem colonization by environmental (e.g., soil) clostridia. For example, for one of our identified acBins (“Tepos_35”), the authors of the original study had also sequenced DNA from soil samples near the burial site (Vågene et al., 2018). *C. tetani* DNA was detected not only in the Tepos_35 sample, but also in the nearby soil although at lower relative abundance (Vågene et al., 2018). Ultimately, this is consistent with the possibility that environmental *C. tetani* from soil may in some cases be the source of the *C. tetani* colonization of ancient human remains. This explanation may account for the observation of low *C. tetani* damage rates in some samples. For other samples where *C. tetani* damage levels are high (>10%) and thus indicative of an ancient origin, it remains unknown whether these strains are the result of ancient sample colonization by environmental strains, or whether they are as old as the archaeological samples themselves.

Regardless of whether the identified *C. tetani* genomes are contemporaneous with the archaeological samples, an important finding of this work is the substantial expansion of genomic knowledge surrounding *C. tetani* and its relatives, such as the expansion of clade 2 and clade 1H, as well as the discovery of *Clostridium* sp. X and Y. Lineage 1H in particular has undergone the greatest expansion through the newly identified aDNA-associated *C. tetani* genomes, from one known sample derived from a patient in France in 2016 (Chapeton-Montes et al., 2019), to 9 additional draft genomes assembled from ancient DNA. This suggests that a broader diversity of 1H strains may exist in undersampled environments. Interestingly, these newly identified lineage 1H strains share a common pattern of originating from the Americas, which suggests that a lineage 1H *C. tetani* strain specific to (or abundant within) this region may have colonized these samples at some point in the past.

In addition to the expansion of existing lineages, our genomic analysis revealed two unique species of *Clostridium* that are closely related to, but distinct from, *C. tetani*. One of these novel species (*Clostridium* sp. “Y”) was assembled from an aDNA sample (GranCanaria-008-Tooth) taken from an archeological specimen dated to 936CE. The other newly discovered species is *Clostridium* sp. “X”, a group of closely related *Clostridium* strains that formed a sister lineage to *C. tetani* and yet are genomically distinct from *C. tetani* and all other available *Clostridium* genomes in the NCBI database. This clade is unlikely to have arisen by errors in genome sequencing or assembly artifacts as it is supported by the co-clustering of multiple genomes, its members are consistently placed as a divergent branch in marker gene phylogenies, its members show a consistent poor coverage alignment to the E88 reference chromosome, and it displays unique gene content and genomic characteristics that are conserved across multiple strains.

Although there appears to be some partial conservation of a *C. tetani* like plasmid in *Clostridium* sp. X and Y with genes such as *repA* and *colT* consistently detected, the *tent* gene was either not detected or was fragmented indicative of a pseudogenization event. These findings support the delineation of the species boundary between these groups, and highlight *tent* as a key gene separating *C. tetani* from closely related species. However, even if they are only pseudogenes, the detection of partial *tent* fragments in the X acBins raises the intriguing possibility that some members of *Clostridium* sp. X may have at one point encoded a full-length *tent* gene, and that *tent*-carrying plasmids may have been circulating in the past outside of *C. tetani* in other closely related *Clostridium* species. This is in line with recent genomic studies that have demonstrated the potential for species outside of the *Clostridium* genus to carry neurotoxin genes (Mansfield et al., 2015; Mansfield & Doxey, 2018; Zhang et al., 2018). Future efforts to sequence the microbiomes associated with archaeological samples, as well as environmental *Clostridium* isolates, may shed further light on these questions.

Beyond expanding *C. tetani* and *Clostridium* genomic diversity, our work also expands the known diversity of clostridial neurotoxins - the most potent family of toxins known to science. Analysis of ancient DNA revealed novel variants and lineages of TeNT, including the newly identified “subgroup 3” toxins: TeNT/Chinchorro, TeNT/El-Yaral toxins, and TeNT/Chiribaya-Alta. Not only do these toxins share a similar mutational profile, but they are derived from a similar geographic area (regions of Peru and Chile in South America) and their associated *C. tetani* genomes also cluster phylogenetically as the closest neighbors. Of the three subgroup 3 *tent* sequences identified, one of them (*tent*/Chinchorro) had sufficient coverage to be fully assembled, and the *tent*/Chinchorro gene also happened to be most divergent from modern *tent* sequences by possessing the greatest number of unique substitutions. Despite being the most divergent *tent*, reads mapping to the *tent*/Chinchorro gene as well as the associated acBin did not show strong patterns of DNA damage, and the damage level was weaker than that for human mtDNA. This indicates that, despite originating from the oldest sample in our dataset and possessing a unique *tent* variant, it is possible that the Chinchorro mummy associated *C. tetani* strain may be a relatively “newer” strain that colonized the sample post-mortem.

Due to the uniqueness of TeNT/Chinchorro, and its collection of amino acid substitutions that have not been observed in any modern TeNT variants, we sought to determine whether this TeNT variant is a functional neurotoxin. Lack of toxicity, for instance, might indicate a sequence artifact or even a TeNT variant that targets other non-mammalian species. We therefore utilized a previous approach based on sortase-mediated ligation to produce small quantities of the fulllength protein toxin (Zhang et al., 2017, 2018). TeNT/Chinchorro produced a classic tetanus phenotype in mouse assays, and exhibits a potency comparable to modern TeNT. This validated our predicted neurotoxic activity for this gene sequence, and suggests that TeNT/Chinchorro’s multiple unique amino acid substitutions have a limited impact on potency and neurotoxicity. However, their non-random spatial clustering on a specific surface-exposed loop of the neurotoxin structure suggests that they may be a result of positive selection, a pattern that has been commonly observed in protein evolutionary studies (Adams et al., 2017; Wagner, 2007). Such substitutions may alter yet-to-be identified TeNT protein-protein interactions. In addition to validating the predicted activity of TeNT/Chinchorro, this experimental method allowed us to directly test the disease-causing properties of a novel virulence factor variant reconstructed from aDNA, without the necessity for growing the organism itself, which would have biosafety concerns.

In summary, using large-scale data mining, we identified ancient neurotoxigenic clostridia in archeological samples. This resulted in a substantial expansion of the known genomic diversity and occurrence of *C. tetani* and led to the discovery of novel *C. tetani* lineages, *Clostridium* species, and tetanus neurotoxin variants that retain functional activity. The discovery of neurotoxigenic clostridial genomes in such a wide diversity of ancient samples, both geographically and temporally, is unexpected, but perhaps not inconsistent with prior hypotheses about the role of these organisms and other toxigenic pathogens in the natural decomposition process (Lobb et al., 2020; Mansfield & Doxey, 2018; Montecucco & Rasotto, 2015). More generally, *Clostridium*-derived DNA is common in ancient DNA collected from human tissues including but not limited to paleofecal microbiomes (Arning & Wilson, 2020) as well as in microbial communities associated with vertebrate decomposition (Javan et al., 2016, 2017).

Despite our identification of neurotoxigenic *C. tetani* and related species in aDNA samples, there are several limitations of our study. These include the limitation of only analyzing the top candidate samples identified from the NCBI-SRA and issues with taxonomic binning and genome assembly of ancient metagenomes. In particular, the main technical limitation of our study is our inability to determine the extent that the identified *C. tetani* and related genomes are derived from one or multiple strains. This leads to some uncertainty in the phylogenetic trees (and estimated damage levels) that we calculated because the patterns we observed may result from a mixture of signals coming from different strains. Lastly, although the damage levels of the identified *C. tetani* and related *Clostridium* genomes are indicative of ancient DNA, the precise origin of these strains remains unknown for the reasons described earlier. Despite this, we anticipate that further exploration of ancient archaeological samples will shed further light on the genomic and functional diversity of these fascinating organisms, as well as the ecology and evolutionary origins of their remarkably potent neurotoxins.

## STAR METHODS

### Identification of *C. tetani* containing samples from the NCBI sequence read archive

To identify datasets within the NCBI sequence read archive predicted to contain *C. tetani* DNA, we searched the NCBI-stat database (Katz et al., 2021) on March 15, 2021 for matches to tax_id=1513 using Google’s Big Query API. The query returned 43,620 sample hits with a *k*-mer self count ranging from 1 to 17,152,980. A threshold of 23,000 was applied which returned 136 hits including 28 *C. tetani* sequencing projects (which were used as controls). The STAT taxonomic profile was retrieved using a separate Google Big Query search and was used to assess the microbial community in each sample. Total counts of each mapped bacterial and archaeal taxon at the species level were extracted from the profile and were converted to proportional values and subsequently visualized in R v4.0.4.

FASTQ files of identified sequencing runs were downloaded to Digital Research Alliance of Canada (formerly Compute Canada) infrastructure using the sra-toolkit v2.9.6, and quality encodings of all runs were assessed. Eight runs were Phred+64 encoded and were converted to Phred+33 using seqtk v1.3 (Shen et al., 2016). Twenty-three runs had an unknown encoding and were assumed to be Phred+33 encoded based on the range of the quality scores. The remaining runs were Phred+33 encoded.

### Recovery and QC assessment of *C. tetani* related genomes from ancient DNA samples

Reads were pre-processed using fastp v0.20.1 (Chen et al., 2018) with default settings to perform quality filtering and remove potential adapters. Pre- and post-processing statistics are included in Table S17. Metagenome co-assembly, using all reads with the same BioSample ID, was performed using MEGAHIT v1.2.9 with default parameters (Li et al., 2014). Contigs were then taxonomically classified using Kaiju v1.7.4 (Menzel et al., 2016) against the Kaiju database nr 2021-02-24 with default settings. Any contigs that mapped to *C. tetani* (NCBI taxonomy ID 1513) or any of its strains (NCBI taxonomy IDs 1231072, 212717, 1172202, and 1172203) were selected for further analyses. The kaiju-identified *C. tetani* contigs were further screened via a BLASTN search with Blast+ v2.12.0 against all *Clostridium* and *Yersinia* species’ genomes from Refseq on Aug. 25, 2022. Any contigs with a *C. tetani* top hit were labeled as *C. tetani* BLAST validated. Contigs containing rRNA, tmRNA, and tRNA were also removed from acBins prior to phylogenetic analysis, as these sequences were associated with high coverage outlier regions due to increased mapping of related reads from non *C. tetani* species. Barrnap v0.9 was used with (--reject 0.05) to sensitively identify rRNA sequences in the metagenomic contigs, including hits up to and including 5% of the expected length. Aragorn v1.2.36 (Laslett & Canback, 2004) was used to identify tRNA and tmRNA sequences with -ps to slightly lower scoring thresholds (to 95% of default).

For QC analysis, CheckM v1.0.18 (Parks et al., 2015) was used to assess acBins for completeness, contamination, and strain heterogeneity, with the pre-built set of *Clostridium* markers supplied with the tool. To examine strain variation, we used a custom Python script to analyze the mpileup results from samtools v1.15.1 (Li et al., 2009) to quantify the mean per-base heterogeneity for each acBin. This involved calculating the percentage of bases for a position in disagreement with the reference base and calculating the overall mean across all non-zero coverage positions.

ANI calculations were performed using two approaches: one method used fastANI v1.33 (Jain et al., 2018) to compare the contig sets for each acBin with *C. tetani* and other *Clostridium* genomes; the second method calculated ANI directly from the snippy-generated multiple genome alignment using the dist.alignment() function from the seqinr v4.2-16 R package (Charif & Lobry, 2007). Scripts associated with the above analyses and contig sets and for three stages of the analysis pipeline (Kaiju, Kaiju+BLAST, Kaiju+BLAST+RNA-removed) are available in the public github repository.

### Measurement and visualization of genome coverage and alignments

Genome coverage for the recovered acBins was further assessed and visualized in two ways. First, Bowtie2 v2.4.2 (Langmead & Salzberg, 2012) was used to map reads from individual runs to the E88 chromosome (accession NC_004557.1) and plasmid (accession NC_004565.1). Bowtie2 was run with the following parameters (--local -D 20 -R 3 -N 1 -L 20 -i S,1,0.50 -bSF4). Using samtools v1.12, the resultant BAM files were then sorted, indexed, and merged based on their BioSample ID. Total (average # of reads per base) and percent (number of bases with 1 or more reads divided by total number of bases) coverage was calculated for the entire chromosome and plasmid as well as the *tent*, *colT*, and *repA* regions. Coverage was visualized using Python v3.8.5 and matplotlib v3.3.2. Coverage plots (Figure S16) were created by loading BAM files into R v4.1.0 with the Rsamtools library v2.8.0 and plotted as area plots using ggplot2 v3.3.5. Coverage of the plasmid sequences was calculated by averaging the number of reads per base in 100bp bins.

As a second approach, we also visualized the multiple genome alignment generated by snippy (described below) using ggplot2 v3.3.5 after loading it into R v4.1.0 using Biostrings 2.62.0.

### Analysis of ancient DNA damage

Fastq files were pre-processed using leeHom v1.2.15 (Renaud et al., 2014) to remove adapters and to perform Bayesian reconstruction of aDNA with the (--ancientdna) flag applied to paired end datasets. The leeHom output was then merged by bioSample ID (concatenated sequentially into one file). Individual and merged results were then processed using seqtk v1.3 with the

“seqtk seq -L30” command to remove short sequences < 30 bp in length. For each BioSample, trimmed reads were then mapped using bwa v0.7.17 (H. Li & Durbin, 2009) to the contigs that classified as *C. tetani* using Kaiju (Menzel et al., 2016) and BLAST with rRNAs, tRNAs, and tmRNAs removed as described above. Trimmed reads were separately mapped to the human mitochondrial reference genome (accession NC_012920.1). Read alignment was performed using “bwa aln” with the (-n 0.01 -o 2 -l 16500) options. BAM files were sorted using samtools v1.12. Misincorporation rates were then measured in two ways: 1) for all samples using mapDamage v2.2.1 (Jónsson et al., 2013) with (--merge-reference-sequences --no-stats) 2) using pyDamage v0.70 (Borry et al., 2021) to estimate DNA damage on a per contig basis with default parameters and per assembly basis with the (--group) parameter.

### Whole genome alignments and phylogenetic reconstruction

All available *C. tetani* genomes (N = 43) were downloaded from the NCBI Genbank database. After duplicate strains were removed, these included the 37 genomes from Chapeton-Montes et al. (2019) as well as four additional genomes that were not included in Chapeton-Montes et al. (2019) but were present in the NCBI (Table S18). To compare our 38 acBins with the 41 non-redundant *C. tetani* genomes and investigate their phylogenetic relationships, we used two approaches described below.

Single base substitutions within assembled *C. tetani* contigs were identified using snippy-multi from the Snippy package v4.6.0 (https://github.com/tseemann/snippy) with *C. tetani* str. E88 as the reference genome (GCA_000007625.1_ASM762v1_genomic.gbff). A genome-wide core alignment was constructed using snippy-core. Five aDNA samples (Deir-Rifeh-KNIII-Tooth, Deir-Rifeh-KNII-Tooth, Pericues-BC28-Bone, Tepos_35-Tooth, Barcelona-3031-Tooth) were removed due to very poor alignment coverage (<1%). Using the resulting alignment, we built a phylogeny using FastTree (Price et al., 2010) v2.1.10 with the GTR model and aLRT metric for assessment of clade support. A maximum-likelihood phylogeny was also constructed using RAxML (Stamatakis, 2014) with a GTR+GAMMA substitution model and 1000 rapid bootstrap inferences. The alignment and phylogeny were analyzed for potential recombination with Gubbins v3.3 (Croucher et al., 2015) and visualized with Phandango v.1.3.0 (Hadfield et al., 2018). The whole genome alignment and trees can be found in the public github repository.

We also used Harvest suite, which provides an automated pipeline for generating and visualizing core-genome alignments, SNP detection, and phylogenetic analysis of intraspecific microbial strains (Treangen et al., 2014). Parsnp v1.2 with the recombination filtration option (-x) was used to examine all acBins and modern *C. tetani* genomes and produce core alignments of orthologous sequences. Since Parsnp only identifies core blocks shared across a group of highly similar genomes, it effectively filters out low coverage and divergent genomes. All 41 modern strains but only 11 acBin strains were kept in the final Parsnp core alignment. FastTree (Price et al., 2010), as included within the Harvest pipeline, was used to produce the final phylogeny. Gingr v1.3 was used to visualize the alignment variation pattern and tree. The resulting tree can be found in the public github repository.

### Annotation and comparative genomics of acBins with modern *C. tetani* strains

We annotated all contigs from the 38 acBins strains using prodigal v2.6.2 (Hyatt et al., 2010). Anonymous annotation mode was used as it is intended for metagenomes and low-quality draft genomes. Sequences encoding rRNA, tRNA and tmRNAs were also annotated as described above. These annotations resulted in predicted proteomes for all 38 acBins, which were then compared with the existing proteomes from the 41 modern *C. tetani* genomes. To cluster identified protein coding sequences into groups of orthologous sequence clusters, we used Orthofinder v2.5.4 (Emms & Kelly, 2019) with default settings. Orthofinder was applied to modern *Clostridium tetani* proteomes and prodigal-predicted acBin proteomes, resulting in a set of 4,334 total orthogroups. Any orthogroup that was made up of only partial coding sequences (as predicted by prodigal) was removed to limit orthogroups resulting from gene fragments or pseudogenes. Orthogroups without any coding sequences found on *C. tetani* BLAST-validated contigs were also removed as a filter for any potential bin contamination. We then performed a gene set comparison between acBins and modern strains by comparing the presence/absence of all identified orthogroups. This resulted in the identification of common orthogroups shared between modern and acBin strains as well as orthogroups unique to acBins, which were investigated further through gene neighborhood analysis. In particular, we focused on orthogroups unique to *Clostridium* sp. X and investigated their gene neighborhoods and their conservation with modern reference *C. tetani* genomes. For gene neighborhood visualization and alignment, we used AnnoView which is located at annoview.uwaterloo.ca.

### Sequence, structural, and phylogenetic analysis of ancient tetanus neurotoxins

*Variant calling and construction of the MSA:* Scripts used for variant calling and generation of a *tent* multiple alignment are located in the public github repository. For the plasmid read alignments used earlier, we extracted aligned reads, and re-aligned them using BWA mem v0.7.17-r1188 using default parameters. Read alignments were manipulated with samtools v1.12 and htslib v1.12. The read alignment was restricted to the *tent* gene locus for variant calling (using the reverse complement of NC_004565.1, bases 1496-5443). Variants were called on each individual sample using the Octopus variant caller v0.7.4 (Cooke et al., 2021) with stringent parameters (--mask-low-quality-tails 5 --min-mapping-quality 10 --min-variant posterior 0.95 -- min-pileup-base-quality 35 --min-good-base-fraction 0.75). This combination of parameters reports only variants with very high confidence and read mapping quality, minimizing identification of false positive variant calls. We then built consensus sequences of *tent* genes from each sample using the bcftools consensus tool v1.12, and htslib v1.12, replacing positions with 0 coverage with a gap character. MAFFT v7.4.80 (Katoh & Standley, 2013) was used to realign fragments against the reference sequence using the (--keeplength) option, which notably keeps the length of the reference unchanged and therefore ignores the possibility of unique insertions. The final *tent* alignments are available in the public github repository.

#### Structural modeling

A structural model of TeNT/Chinchorro was generated by automated homology modeling using the SWISSMODEL server (Waterhouse et al., 2018). Modeling was performed using two top-scoring homologous template structures of tetanus neurotoxins: PDB IDs 7by5.1.A (97.18% identity), 5N0C.1.A (97.34% identity). 7BY5.1.A was selected as the best template based on the QMEAN quality estimate (Benkert et al., 2011). The model was visualized using PyMOL v2.4.1 and unique substitutions (present in TeNT/Chinchorro but absent in modern TeNT sequences) were highlighted.

#### Phylogenetic analysis

The *tent* consensus alignment generated as described earlier was processed to keep only sequences (N = 20) with alignment coverage exceeding 80%. The following BioSamples were removed: Tepos_35-Tooth, Tenerife-012-Tooth, Tenerife-011-Tooth, SLC-France-Tooth, Peru-NA42-Bone, Deir-Rifeh-KNII-Tooth, Peru-NA40-Bone, Punta_Candelero-Tooth, Gniezno-Tooth, Augsburg-Tooth, Riga_Cemetery-Tooth, Pericues-BC28-Bone, Marseille-Tooth. We then aligned the 20 ancient *tent* gene sequences with 30 *tent* sequences from modern *C. tetani* strains, which reduced to 12 representative modern *tent* sequences after duplicates were removed using Jalview v2.9.0b2. *tent/E88* was identical with *tent* from 11 strains (1586-U1, CN655, 641.84, C2, Strain_3, 75.97, 89.12, 46.1.08, A, 4784A, Harvard), *tent*/132CV with 1 other (Mfbjulcb2), *tent*/63.05 with 2 others (3483, 184.08), *tent*/1337 with 2 others (B4, 1240), *tent*/ATCC_453 with 1 other (3582), and *tent*/202.15 with 1 other (358.99). A phylogeny was constructed using PhyML v3.1 (Guindon & Gascuel, 2003) with the GTR model, empirical nucleotide equilibrium frequencies, no invariable sites, across site rate variation optimized, NNI tree search, and BioNJ as the starting tree. PhyML analysis identified 362 patterns, and aLRT (SH-like) branch supports were calculated. The final newick tree is available in the public github repository.

### Experimental testing of TeNT/Chinochorro (chTeNT)

*Antibodies and constructs:* Antibodies for Syntaxin-1 (HPC-1), SNAP25 (C171.2), VAMP1/2/3 (104102) were purchased from Synaptic Systems. Antibody against actin (AC-15) was purchased from Sigma. Rabbit antiserum of TeNT (ab53829) was purchased from Abcam, rabbit nonimmunized serum (AB110) was purchased from Boston Molecules. The cDNAs encoding chTeNT-LC-H_N_ (the N-terminal fragment, residues 1-870) and chTeNT-H_C_ (the C-terminal fragment, residues 875-1315) were synthesized by Twist Bioscience (South San Francisco, CA). The cDNA encoding TeNT-LC-H_N_ (residues 1-870) and TeNT-H_C_ were synthesized by GenScript (Piscataway, NJ). A thrombin protease cleavage site was inserted between I448 and A457 in both TeNT-LC-H_N_ and chTeNT-LC-H_N_. LC-H_N_ fragments were cloned into pET28a vector, with peptide sequence LPETGG fused to their C-termini, followed by a His6-tag. H_C_ fragments were cloned into pET28a vectors with a His6-tag and thrombin recognition site on their N-termini.

#### Protein purification

*E. coli* BL21 (DE3) was utilized for protein expression. In general, transformed bacteria were cultured in LB medium using an orbital shaker at 37 °C until the OD600 reached 0.6. Induction of protein expression was carried out with 0.1 mM IPTG at 18 °C overnight. Bacterial pellets were collected by centrifugation at 4,000 g for 10 min and disrupted by sonication in lysis buffer (50 mM Tris pH 7.5, 250 mM NaCl, 1 mM PMSF, 0.4 mM lysozyme), and supernatants were collected after centrifugation at 20,000 g for 30 min at 4 °C. Protein purification was carried out using a gravity nickel column, then purified proteins were desalted with PD-10 columns (GE, 17-0851-01) and concentrated using Centrifugal Filter Units (EMD Millipore, UFC803008).

#### Sortase ligation

H_C_ protein fragments were cleaved by thrombin (40 mU/μL) (EMD Millipore, 605157-1KU) overnight at 4 °C. Ligation reaction was set up in 100 μL TBS buffer with LC-H_N_ (8 μM), H_C_ (5 μM), Ca2+ (10mM) and sortase (1.5 μM), for 1 hour at room temperature. Full-length proteins were then activated by thrombin (40 mU/μL) at room temperature for 1 hour. Sortase ligation reaction mixtures were analyzed by Coomassie blue staining and quantified by BSA reference standards.

#### Neuron culture and immunoblot analysis

Primary rat cortical neurons were prepared from E18-19 embryos using a papain dissociation kit (Worthington Biochemical) following the manufacturer’s instruction. Neurons were exposed to sortase ligation mixtures with or without antiserum in culture medium for 12 hrs. Cells were then lysed with RIPA buffer with protease inhibitor cocktail (Sigma-Aldrich). Lysates were centrifuged at 12000 g at 4 °C for 10 min. Supernatants were subjected to SDS–PAGE and immunoblot analysis.

#### Animal study

All animal studies were approved by the Boston Children’s Hospital Institutional Animal Care and Use Committee (Protocol Number: 18-10-3794R). Toxins were diluted using phosphate buffer (pH 6.3) containing 0.2% gelatin. Mice (CD-1 strain, female, purchased from Envigo, 6-7 weeks old, 25–28 g, n=3 per group, 9 total) were anesthetized with isoflurane (3–4%) and injected with toxin (10 μL) using a 30-gauge needle attached to a sterile Hamilton syringe, into the gastrocnemius muscles of the right hind limb with the left leg serving as the negative control. Muscle paralysis was observed for 4 days. The severity of spastic paralysis was scored with a numerical scale modified from a previous report (0, no symptoms; 4, injected limb and toes are fully rigid) (Mellanby et al., 1968).

## ACKNOWLEDGEMENTS

This study was supported by the Natural Sciences and Engineering Research Council (NSERC) through a Discovery Grant (RGPIN-2019-04266) and Discovery Accelerator Supplement (RGPAS-2019-00004) awarded to A.C.D., by the Government of Ontario through an Early Researcher Award to A.C.D, and by a University of Waterloo Interdisciplinary Trailblazer grant awarded to A.C.D. and A.D. A.C.D. also holds a University Research Chair from the University of Waterloo. M.J.M. gratefully acknowledges funding from the Japan Society for the Promotion of Science as a JSPS International Research Fellow (Luscombe Unit, Okinawa Institute of Science and Technology Graduate University). H.P.H. gratefully acknowledges funding from an NSERC Canada Graduate Scholarship. H.P.H. and A.C.D. also acknowledge the Digital Research Alliance of Canada (formerly Compute Canada) for providing access to high performance computing resources. This study was also partially supported by grants from National Institute of Health (NIH) (R01NS080833 and R01NS117626 to M.D). M.D. holds the Investigator in the Pathogenesis of Infectious Disease award from the Burroughs Wellcome Fund.

## AUTHOR CONTRIBUTIONS

A.C.D. conceived and supervised the project. H.P.H., B.T., B.L., M.J.M, V.L., X.W., and A.C.D. performed bioinformatic data analysis. A.C.D. and G.R. supervised aDNA analysis. P.C. and P.L. performed all experimental work, which was supervised by M.D. A.D. and J.C. performed context analysis of archaeological samples. All authors contributed to manuscript writing and preparation of figures.

## DECLARATION OF INTERESTS

The authors declare no competing interests.

## DATA AND CODE AVAILABILITY

Genomic data reported in this study is available from the NCBI sequence read archive. Accession numbers for all BioSamples and sequencing runs used are listed in the Supplemental Information. Source code and additional data for this project are available at https://github.com/harohodg/aDNA-tetanus-analysis

## Notes

### Competing Interest Statement

The authors have declared no competing interest.

### Summary of Updates

Expanded genomic analysis of "X" and "Y" strains; updated analysis of DNA damage levels after subdividing datasets by UDG treatment and capture methods; additional phylogenetic and recombination analysis; added genomic analysis of regions surrounding key genes of interest; additional experimental data on newly identified toxin; comprehensive QC analysis of bins

## REFERENCES

Adams, J., Mansfield, M. J., Richard, D. J., & Doxey, A. C. (2017). Lineage-specific mutational clustering in protein structures predicts evolutionary shifts in function. Bioinformatics, 33(9), 1338–1345.

Adler, C. J., Haak, W., Donlon, D., & Cooper, A. (2011). Survival and recovery of DNA from ancient teeth and bones. Journal of Archaeological Science, 38(5), 956–964.

Altschul, S. F., Madden, T. L., Schäffer, A. A., Zhang, J., Zhang, Z., Miller, W., & Lipman, D. J. (1997). Gapped BLAST and PSI-BLAST: A new generation of protein database search programs. Nucleic Acids Research, 25, 3389–3402.

Andrades Valtueña, A., Mittnik, A., Key, F. M., Haak, W., Allmäe, R., Belinskij, A., Daubaras, M., Feldman, M., Jankauskas, R., Janković, I., Massy, K., Novak, M., Pfrengle, S., Reinhold, S., Šlaus, M., Spyrou, M. A., Szécsényi-Nagy, A., Tõrv, M., Hansen, S.,… Krause, J. (2017). The Stone Age Plague and Its Persistence in Eurasia. Current Biology: CB, 27(23), 3683–3691.e8.

Arning, N., & Wilson, D. J. (2020). The past, present and future of ancient bacterial DNA. Microbial Genomics, 6(7), 1–19.

Benkert, P., Biasini, M., & Schwede, T. (2011). Toward the estimation of the absolute quality of individual protein structure models. Bioinformatics (Oxford, England), 27(3), 343–350.

Borry, M., Hübner, A., Rohrlach, A. B., & Warinner, C. (2021). PyDamage: automated ancient damage identification and estimation for contigs in ancient DNA de novo assembly. PeerJ, 9.

Bos, K., Harkins, K., Herbig, A., Coscolla, M., Weber, N., Comas, I., Forrest, S., Bryant, J., Harris, S., Schuenemann, V., Campbell, T., Majander, K., Wilbur, A., Guichon, R., Wolfe Steadman, D., Cook, D., Niemann, S., Behr, M., Zumarraga, M.,… Krause, J. (2014). Pre-Columbian mycobacterial genomes reveal seals as a source of New World human tuberculosis. Nature, 514(7523), 494–497.

Bos, K. I., Herbig, A., Sahl, J., Waglechner, N., Fourment, M., Forrest, S. A., Klunk, J., Schuenemann, V. J., Poinar, D., Kuch, M., Golding, G. B., Dutour, O., Keim, P., Wagner, D. M., Holmes, E. C., Krause, J., & Poinar, H. N. (2016). Eighteenth century Yersinia pestis genomes reveal the long-term persistence of an historical plague focus. ELife, 5.

Bos, K., Kühnert, D., Herbig, A., Esquivel-Gomez, L., Andrades Valtueña, A., Barquera, R., Giffin, K., Kumar Lankapalli, A., Nelson, E., Sabin, S., Spyrou, M., & Krause, J. (2019). Paleomicrobiology: Diagnosis and Evolution of Ancient Pathogens. Annual Review of Microbiology, 73, 639–666.

Bos, K., Schuenemann, V., Golding, G., Burbano, H., Waglechner, N., Coombes, B., McPhee, J., DeWitte, S., Meyer, M., Schmedes, S., Wood, J., Earn, D., Herring, D., Bauer, P., Poinar, H., & Krause, J. (2011). A draft genome of Yersinia pestis from victims of the Black Death. Nature, 478(7370), 506–510.

Briggs, A., Stenzel, U., Johnson, P., Green, R., Kelso, J., Prüfer, K., Meyer, M., Krause, J., Ronan, M., Lachmann, M., & Pääbo, S. (2007). Patterns of damage in genomic DNA sequences from a Neandertal. Proceedings of the National Academy of Sciences of the United States of America, 104(37), 14616–14621.

Bruggemann, H., Baumer, S., Fricke, W., Wiezer, A., Liesegang, H., Decker, I., Herzberg, C., Martinez-Arias, R., Merkl, R., Henne, A., & Gottschalk, G. (2003). The genome sequence of Clostridium tetani, the causative agent of tetanus disease. Proceedings of the National Academy of Sciences of the United States of America, 100(3), 1316–1321.

Brüggemann, H., Brzuszkiewicz, E., Chapeton-Montes, D., Plourde, L., Speck, D., & Popoff, M. R. (2015). Genomics of Clostridium tetani. Research in Microbiology, 166(4), 326–331.

Chapeton-Montes, D., Plourde, L., Bouchier, C., Ma, L., Diancourt, L., Criscuolo, A., Popoff, M., & Brüggemann, H. (2019). The population structure of Clostridium tetani deduced from its pan-genome. Scientific Reports, 9(1).

Charif, D., & Lobry, J. R. (2007). SeqinR 1.0-2: A Contributed Package to the R Project for Statistical Computing Devoted to Biological Sequences Retrieval and Analysis (pp. 207–232). Springer, Berlin, Heidelberg.

Chen, S., Zhou, Y., Chen, Y., & Gu, J. (2018). fastp: an ultra-fast all-in-one FASTQ preprocessor. Bioinformatics (Oxford, England), 34(17), i884–i890.

Cohen, J. E., Wang, R., Shen, R. F., Wu, W. W., & Keller, J. E. (2017). Comparative pathogenomics of Clostridium tetani. PLoS ONE, 12(8).

Cooke, D., Wedge, D., & Lunter, G. (2021). A unified haplotype-based method for accurate and comprehensive variant calling. Nature Biotechnology, 39(7), 885–892.

Croucher, N. J., Page, A. J., Connor, T. R., Delaney, A. J., Keane, J. A., Bentley, S. D., Parkhill, J., & Harris, S. R. (2015). Rapid phylogenetic analysis of large samples of recombinant bacterial whole genome sequences using Gubbins. Nucleic Acids Research, 43(3), e15.

Dabney, J., Meyer, M., & Pääbo, S. (2013). Ancient DNA damage. Cold Spring Harbor Perspectives in Biology, 5(7).

Darraj, M., Stone, J., Keynan, Y., Thompson, K., & Snider, C. (2017). A case of tetanus secondary to an odontogenic infection. CJEM, 19(6), 497–499.

de la Fuente, C., Ávila-Arcos, M. C., Galimany, J., Carpenter, M. L., Homburger, J. R., Blanco, A., Contreras, P., Dávalos, D. C., Reyes, O., Roman, M. S., Moreno-Estrada, A., Campos, P. F., Eng, C., Huntsman, S., Burchard, E. G., Malaspinas, A. S., Bustamante, C. D., Willerslev, E., Llop, E.,… Moraga, M. (2018). Genomic insights into the origin and diversification of late maritime hunter-gatherers from the Chilean Patagonia. Proceedings of the National Academy of Sciences of the United States of America, 115(17), E4006–E4012.

Drancourt, M., & Raoult, D. (2005). Palaeomicrobiology: current issues and perspectives. Nature Reviews. Microbiology, 3(1), 23–35.

Duggan, A. T., Perdomo, M. F., Piombino-Mascali, D., Marciniak, S., Poinar, D., Emery, M. v., Buchmann, J. P., Duchêne, S., Jankauskas, R., Humphreys, M., Golding, G. B., Southon, J., Devault, A., Rouillard, J. M., Sahl, J. W., Dutour, O., Hedman, K., Sajantila, A., Smith, G. L.,… Poinar, H. N. (2016). 17 th Century Variola Virus Reveals the Recent History of Smallpox. Current Biology: CB, 26(24), 3407–3412.

Emms, D. M., & Kelly, S. (2019). OrthoFinder: phylogenetic orthology inference for comparative genomics. Genome Biology, 20(1).

Guindon, S., & Gascuel, O. (2003). A simple, fast, and accurate algorithm to estimate large phylogenies by maximum likelihood. Systematic Biology, 52, 696–704.

Hadfield, J., Croucher, N. J., Goater, R. J., Abudahab, K., Aanensen, D. M., & Harris, S. R. (2018). Phandango: an interactive viewer for bacterial population genomics. Bioinformatics, 34(2), 292.

Hyatt, D., Chen, G.-L., Locascio, P. F., Land, M. L., Larimer, F. W., & Hauser, L. J. (2010). Prodigal: prokaryotic gene recognition and translation initiation site identification. BMC Bioinformatics, 11(1), 119.

Jain, C., Rodriguez-R, L. M., Phillippy, A. M., Konstantinidis, K. T., & Aluru, S. (2018). High throughput ANI analysis of 90K prokaryotic genomes reveals clear species boundaries. Nature Communications, 9(1).

Javan, G. T., Finley, S. J., Can, I., Wilkinson, J. E., Hanson, J. D., & Tarone, A. M. (2016). Human Thanatomicrobiome Succession and Time Since Death. Scientific Reports, 6(1), 29598.

Javan, G. T., Finley, S. J., Smith, T., Miller, J., & Wilkinson, J. E. (2017). Cadaver Thanatomicrobiome Signatures: The Ubiquitous Nature of Clostridium Species in Human Decomposition. Frontiers in Microbiology, 8(OCT).

Jónsson, H., Ginolhac, A., Schubert, M., Johnson, P. L. F., & Orlando, L. (2013). mapDamage2.0: fast approximate Bayesian estimates of ancient DNA damage parameters. Bioinformatics (Oxford, England), 29(13), 1682–1684.

Kanzawa-Kiriyama, H., Kryukov, K., Jinam, T., Hosomichi, K., Saso, A., Suwa, G., Ueda, S., Yoneda, M., Tajima, A., Shinoda, K., Inoue, I., & Saitou, N. (2017). A partial nuclear genome of the Jomons who lived 3000 years ago in Fukushima, Japan. Journal of Human Genetics, 62(2), 213–221.

Katoh, K., & Standley, D. M. (2013). MAFFT multiple sequence alignment software version 7: improvements in performance and usability. Molecular Biology and Evolution, 30(4), 772–780.

Katz, K., Shutov, O., Lapoint, R., Kimelman, M., Brister, J., & O’Sullivan, C. (2021). STAT: a fast, scalable, MinHash-based k-mer tool to assess Sequence Read Archive next-generation sequence submissions. Genome Biology, 22(1).

Kitasato, S. (1889). Ueber den Tetanusbacillus. Z Hyg, 7, 225–34.

Langmead, B., & Salzberg, S. L. (2012). Fast gapped-read alignment with Bowtie 2. Nature Methods, 9(4), 357–359.

Laslett, D., & Canback, B. (2004). ARAGORN, a program to detect tRNA genes and tmRNA genes in nucleotide sequences. Nucleic Acids Research, 32(1), 11–16.

Levy, P., Fournier, P., Lotte, L., Million, M., Brouqui, P., & Raoult, D. (2014). Clostridium tetani osteitis without tetanus. Emerging Infectious Diseases, 20(9), 1571–1573.

Li, D., Liu, C. M., Luo, R., Sadakane, K., & Lam, T. W. (2014). MEGAHIT: An ultra-fast single-node solution for large and complex metagenomics assembly via succinct de Bruijn graph. Bioinformatics, 31(10), 1674–1676.

Li, H., & Durbin, R. (2009). Fast and accurate short read alignment with Burrows-Wheeler transform. Bioinformatics, 25(14), 1754–1760.

Li, H., Handsaker, B., Wysoker, A., Fennell, T., Ruan, J., Homer, N., Marth, G., Abecasis, G., & Durbin, R. (2009). The Sequence Alignment/Map format and SAMtools. Bioinformatics (Oxford, England), 25(16), 2078–2079.

Lobb, B., Hodgson, R., Lynch, M. D. J., Mansfield, M. J., Cheng, J., Charles, T. C., Neufeld, J. D., Craig, P. M., & Doxey, A. C. (2020). Time Series Resolution of the Fish Necrobiome Reveals a Decomposer Succession Involving Toxigenic Bacterial Pathogens. MSystems, 5(2).

Maixner, F., Krause-Kyora, B., Turaev, D., Herbig, A., Hoopmann, M. R., Hallows, J. L., Kusebauch, U., Vigl, E. E., Malfertheiner, P., Megraud, F., O’Sullivan, N., Cipollini, G., Coia, V., Samadelli, M., Engstrand, L., Linz, B., Moritz, R. L., Grimm, R., Krause, J.,… Zink, A. (2016). The 5300-year-old Helicobacter pylori genome of the Iceman. Science (New York, N.Y.), 351(6269), 162–165.

Malmström, H., Günther, T., Svensson, E. M., Juras, A., Fraser, M., Munters, A. R., Pospieszny, Ł., Tõrv, M., Lindström, J., Götherström, A., Storå, J., & Jakobsson, M. (2019). The genomic ancestry of the Scandinavian Battle Axe Culture people and their relation to the broader Corded Ware horizon. Proceedings. Biological Sciences, 286(1912).

Mansfield, M. J., Adams, J. B., & Doxey, A. C. (2015). Botulinum neurotoxin homologs in non-Clostridium species. FEBS Letters, 589(3), 342–348.

Mansfield, M. J., & Doxey, A. C. (2018). Genomic insights into the evolution and ecology of botulinum neurotoxins. Pathogens and Disease, 76(4).

Masuyer, G., Conrad, J., & Stenmark, P. (2017). The structure of the tetanus toxin reveals pH-mediated domain dynamics. EMBO Reports, 18(8), 1306–1317.

Megighian, A., Pirazzini, M., Fabris, F., Rossetto, R., & Montecucco, C. (2021). Tetanus and tetanus neurotoxin: From peripheral uptake to central nervous tissue targets. Journal of Neurochemistry, 158(6), 1244–1253.

Mellanby, J., Mellanby, H., Pope, D., & van Heyningen, W. (1968). Ganglioside as a prophylactic agent in experimental tetanus in mice. J Gen Microbiol, 54(2), 161–168.

Mendler, K., Chen, H., Parks, D. H., Lobb, B., Hug, L. A., & Doxey, A. C. (2019). AnnoTree: visualization and exploration of a functionally annotated microbial tree of life. Nucleic Acids Research, 47(9), 4442–4448.

Menzel, P., Ng, K., & Krogh, A. (2016). Fast and sensitive taxonomic classification for metagenomics with Kaiju. Nature Communications, 7.

Miles, S. H. (2009). Hippocrates and informed consent. Lancet (London, England), 374(9698), 1322–1323.

Montecucco, C., & Rasotto, M. B. (2015). On botulinum neurotoxin variability. MBio, 6(1), e02131–14.

Mühlemann, B., Jones, T., Damgaard, P., Allentoft, M., Shevnina, I., Logvin, A., Usmanova, E., Panyushkina, I., Boldgiv, B., Bazartseren, T., Tashbaeva, K., Merz, V., Lau, N., Smrčka, V., Voyakin, D., Kitov, E., Epimakhov, A., Pokutta, D., Vicze, M.,… Willerslev, E. (2018). Ancient hepatitis B viruses from the Bronze Age to the Medieval period. Nature, 557(7705), 418–423.

Namouchi, A., Guellil, M., Kersten, O., Hänsch, S., Ottoni, C., Schmid, B. v., Pacciani, E., Quaglia, L., Vermunt, M., Bauer, E. L., Derrick, M., Jensen, A., Kacki, S., Cohn, S. K., Stenseth, N. C., & Bramanti, B. (2018). Integrative approach using Yersinia pestis genomes to revisit the historical landscape of plague during the Medieval Period. Proceedings of the National Academy of Sciences of the United States of America, 115(50), E11790–E11797.

Neukamm, J., Pfrengle, S., Molak, M., Seitz, A., Francken, M., Eppenberger, P., Avanzi, C., Reiter, E., Urban, C., Welte, B., Stockhammer, P., Teßmann, B., Herbig, A., Harvati, K., Nieselt, K., Krause, J., & Schuenemann, V. (2020). 2000-year-old pathogen genomes reconstructed from metagenomic analysis of Egyptian mummified individuals. BMC Biology, 18(1).

Pappas, G., Kiriaze, I. J., & Falagas, M. E. (2008). Insights into infectious disease in the era of Hippocrates. International Journal of Infectious Diseases: IJID: Official Publication of the International Society for Infectious Diseases, 12(4), 347–350.

Parks, D. H., Chuvochina, M., Waite, D. W., Rinke, C., Skarshewski, A., Chaumeil, P.-A., & Hugenholtz, P. (2018). A standardized bacterial taxonomy based on genome phylogeny substantially revises the tree of life. Nature Biotechnology.

Parks, D., Imelfort, M., Skennerton, C., Hugenholtz, P., & Tyson, G. (2015). CheckM: assessing the quality of microbial genomes recovered from isolates, single cells, and metagenomes. Genome Research, 25(7), 1043–1055.

Popoff, M. (2020). Tetanus in animals. J Vet Diagn Invest, 32(2), 184–191.

Price, M., Dehal, P., & Arkin, A. (2010). FastTree 2--approximately maximum-likelihood trees for large alignments. PloS One, 5(3).

Raghavan, M., Steinrücken, M., Harris, K., Schiffels, S., Rasmussen, S., DeGiorgio, M., Albrechtsen, A., Valdiosera, C., Ávila-Arcos, M., Malaspinas, A., Eriksson, A., Moltke, I., Metspalu, M., Homburger, J., Wall, J., Cornejo, O., Moreno-Mayar, J., Korneliussen, T., Pierre, T.,… Willerslev, E. (2015). Genomic evidence for the Pleistocene and recent population history of Native Americans. Science (New York, N.Y.), 349(6250).

Renaud, G., Stenzel, U., & Kelso, J. (2014). leeHom: adaptor trimming and merging for Illumina sequencing reads. Nucleic Acids Research, 42(18), e141.

Richter, M., & Rosselló-Móra, R. (2009). Shifting the genomic gold standard for the prokaryotic species definition. Proceedings of the National Academy of Sciences of the United States of America, 106(45), 19126–19131.

Rodríguez-Varela, R., Günther, T., Krzewińska, M., Storå, J., Gillingwater, T., MacCallum, M., Arsuaga, J., Dobney, K., Valdiosera, C., Jakobsson, M., Götherström, A., & Girdland-Flink, L. (2017). Genomic Analyses of Pre-European Conquest Human Remains from the Canary Islands Reveal Close Affinity to Modern North Africans. Current Biology: CB, 27(21), 3396–3402.e5.

Rohland, N., Harney, E., Mallick, S., Nordenfelt, S., & Reich, D. (2015). Partial uracil-DNA-glycosylase treatment for screening of ancient DNA. Philosophical Transactions of the Royal Society of London. Series B, Biological Sciences, 370(1660).

Sabin, S., Herbig, A., Vågene, Å. J., Ahlström, T., Bozovic, G., Arcini, C., Kühnert, D., & Bos, K. I. (2020). A seventeenth-century Mycobacterium tuberculosis genome supports a Neolithic emergence of the Mycobacterium tuberculosis complex. Genome Biology, 21(1).

Sanchez, G. M., & Burridge, A. L. (2007). Decision making in head injury management in the Edwin Smith Papyrus. Neurosurgical Focus, 23(1).

Sawyer, S., Krause, J., Guschanski, K., Savolainen, V., & Pääbo, S. (2012). Temporal patterns of nucleotide misincorporations and DNA fragmentation in ancient DNA. PLoS ONE, 7(3).

Schuenemann, V., Singh, P., Mendum, T., Krause-Kyora, B., Jäger, G., Bos, K., Herbig, A., Economou, C., Benjak, A., Busso, P., Nebel, A., Boldsen, J., Kjellström, A., Wu, H., Stewart, G., Taylor, G., Bauer, P., Lee, O., Wu, H.,… Krause, J. (2013). Genome-wide comparison of medieval and modern Mycobacterium leprae. Science (New York, N.Y.), 341(6142), 179–183.

Shen, W., Le, S., Li, Y., & Hu, F. (2016). SeqKit: A Cross-Platform and Ultrafast Toolkit for FASTA/Q File Manipulation. PloS One, 11(10).

Stamatakis, A. (2014). RAxML version 8: a tool for phylogenetic analysis and post-analysis of large phylogenies. Bioinformatics, 30(9), 1312–1313.

Stolarek, I., Juras, A., Handschuh, L., Marcinkowska-Swojak, M., Philips, A., Zenczak, M., Dębski, A., Kóčka-Krenz, H., Piontek, J., Kozlowski, P., & Figlerowicz, M. (2018). A mosaic genetic structure of the human population living in the South Baltic region during the Iron Age. Scientific Reports, 8(1).

Susat, J., Bonczarowska, J. H., Pētersone-Gordina, E., Immel, A., Nebel, A., Gerhards, G., & Krause-Kyora, B. (2020). Yersinia pestis strains from Latvia show depletion of the pla virulence gene at the end of the second plague pandemic. Scientific Reports, 10(1).

Treangen, T. J., Ondov, B. D., Koren, S., & Phillippy, A. M. (2014). The Harvest suite for rapid core-genome alignment and visualization of thousands of intraspecific microbial genomes. Genome Biology, 15(11).

Vågene, Å. J., Herbig, A., Campana, M. G., Robles García, N. M., Warinner, C., Sabin, S., Spyrou, M. A., Andrades Valtueña, A., Huson, D., Tuross, N., Bos, K. I., & Krause, J. (2018). Salmonella enterica genomes from victims of a major sixteenth-century epidemic in Mexico. Nature Ecology & Evolution, 2(3), 520–528.

Valdiosera, C., Günther, T., Vera-Rodríguez, J. C., Ureña, I., Iriarte, E., Rodríguez-Varela, R., Simões, L. G., Martínez-Sánchez, R. M., Svensson, E. M., Malmström, H., Rodríguez, L., de Castro, J. M. B., Carbonell, E., Alday, A., Vera, J. A. H., Götherström, A., Carretero, J. M., Arsuaga, J. L., Smith, C. I., & Jakobsson, M. (2018). Four millennia of Iberian biomolecular prehistory illustrate the impact of prehistoric migrations at the far end of Eurasia. Proceedings of the National Academy of Sciences of the United States of America, 115(13), 3428–3433.

Wagner, A. (2007). Rapid detection of positive selection in genes and genomes through variation clusters. Genetics, 176(4), 2451–2463.

Warinner, C., Speller, C., & Collins, M. (2015). A new era in palaeomicrobiology: prospects for ancient dental calculus as a long-term record of the human oral microbiome. Philosophical Transactions of the Royal Society of London. Series B, Biological Sciences, 370(1660).

Waterhouse, A., Bertoni, M., Bienert, S., Studer, G., Tauriello, G., Gumienny, R., Heer, F., de Beer, T., Rempfer, C., Bordoli, L., Lepore, R., & Schwede, T. (2018). SWISS-MODEL: homology modelling of protein structures and complexes. Nucleic Acids Research, 46(W1), W296–W303.

Zhang, S., Lebreton, F., Mansfield, M. J., Miyashita, S.-I., Zhang, J., Schwartzman, J. A., Tao, L., Masuyer, G., Martínez-Carranza, M., Stenmark, P., Gilmore, M. S., Doxey, A. C., & Dong, M. (2018). Identification of a Botulinum Neurotoxin-like Toxin in a Commensal Strain of Enterococcus faecium. Cell Host & Microbe, 23(2), 169–176.e6.

Zhang, S., Masuyer, G., Zhang, J., Shen, Y., Lundin, D., Henriksson, L., Miyashita, S.-I., Martínez-Carranza, M., Dong, M., & Stenmark, P. (2017). Identification and characterization of a novel botulinum neurotoxin. Nature Communications, 8, 14130.

