## Supplementary Figures for "Ancient *Clostridium* DNA and variants of tetanus neurotoxins associated with human archaeological remains"

### SUPPLEMENTAL INFORMATION

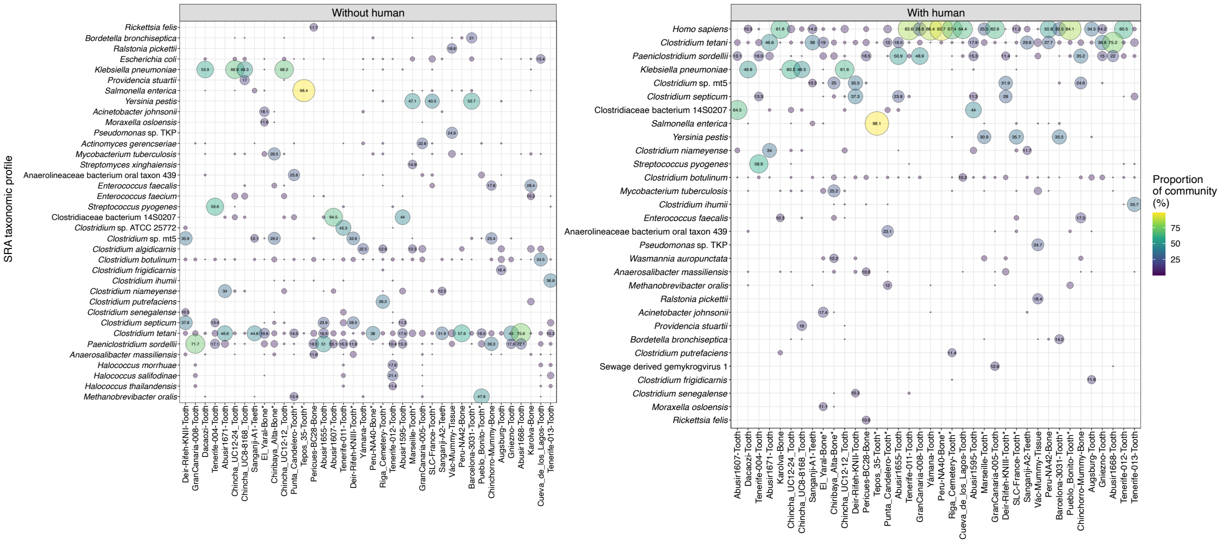


**Figure S1. Proportional abundance of taxa detected in metagenomes from 38 aDNA samples (microbial only – left, all – right).** Abundance values are based on NCBI sequence read archive taxonomic profiles including all identified bacterial and archaeal species. Only species with >10% abundance in at least one sample have been plotted, and the full dataset is available in Table S3. Only values greater than 10% are labelled in the figure. For BioSamples associated with more than one SRA ID, the sample with a median count of *Clostridium tetani* was chosen. In the case of a choice between two, a random SRA ID was chosen. The samples here are represented as follows: Sanganji-A1-Teeth: DRR046398; Sanganji-A2-Teeth: DRR046408; Augsburg-Tooth: ERR2112574; GranCanaria-005-Tooth: ERR2111951; GranCanaria-008-Tooth: ERR2112080; Chinchorro-Mummy-Bone: ERR966303; SLC-France-Tooth: ERR2862150; Peru-NA42-Bone: SRR1298752; Pueblo_Bonito-Tooth; SRR5169887. *Sample names with asterisks indicate those associated with capture sequencing methods.

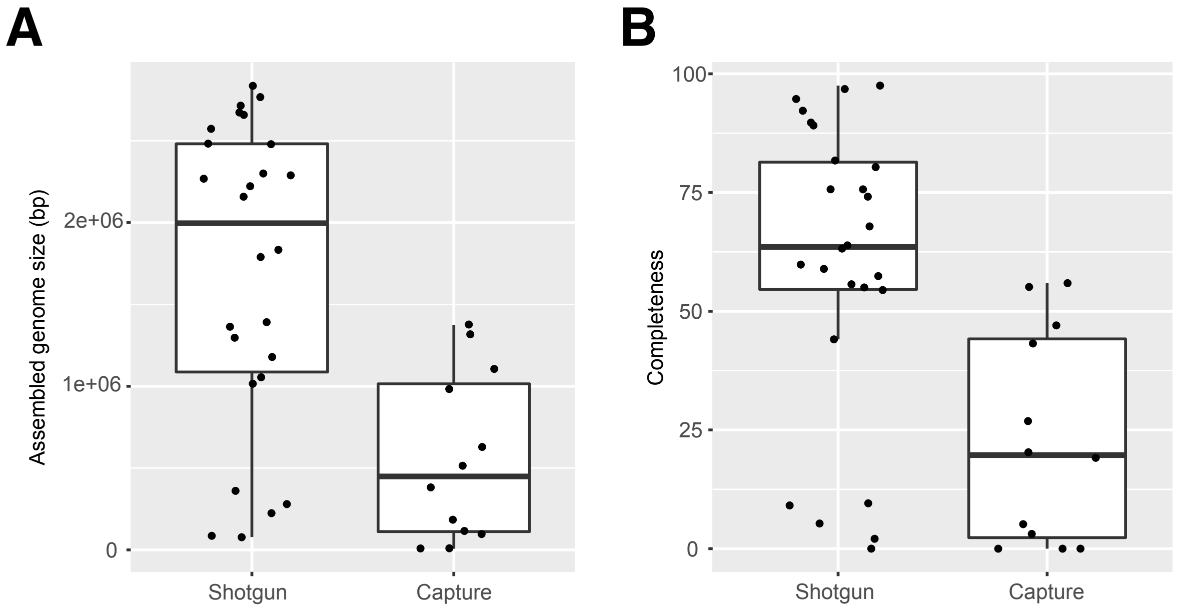


**Figure S2. Assembly size and completeness of recovered acBins based on use of shotgun versus capture sequencing methods.** The assembly size and completeness statistics were calculated using CheckM and are available in Table S6. Assembled genome size and estimated completeness were both higher when capture methods were not used in the aDNA sequencing protocol.

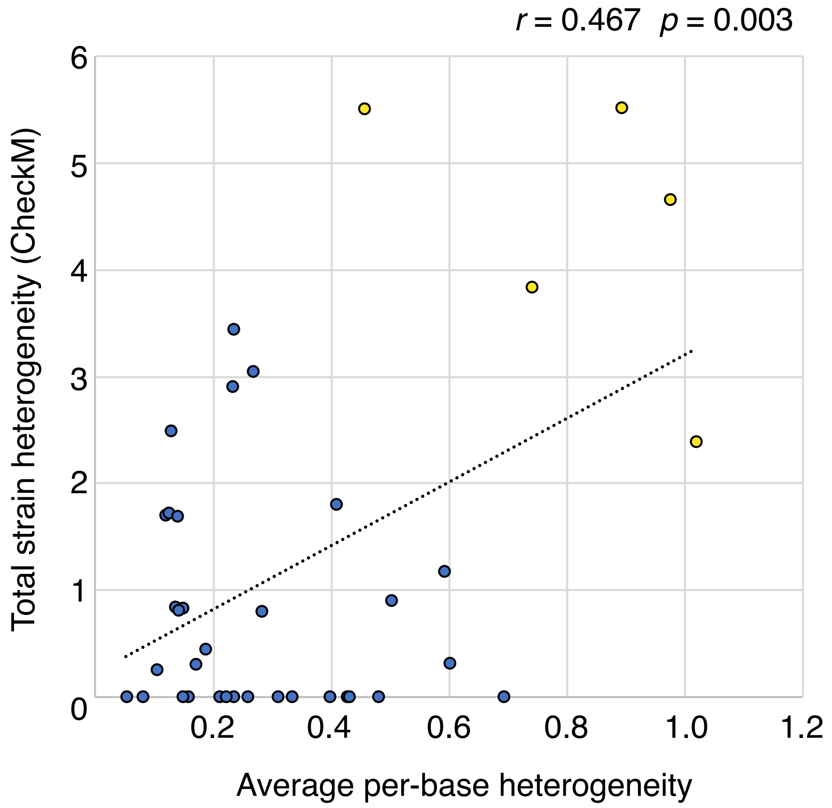


**Figure S3. Comparison of strain heterogeneity levels estimated from mapped base heterogeneity versus CheckM calculations.** Average per-base heterogeneity (x-axis) was calculated for each acBin by examining mapped reads for all positions across contigs, and calculating the fraction of bases that disagree with the reference base. We then calculated the mean of the per-base fractional heterogeneity value across all non-zero coverage positions. Total strain heterogeneity was calculated as the product of contamination and the strain heterogeneity as measured using CheckM. The five samples with the highest probability of containing strain variation are shown as yellow circles. These include: Sanganji-A2-Tooth, Chinchorro-Mummy-Bone, SLC-France-Tooth, Karolva-Tooth, Chincha-UC12-24-Tooth.

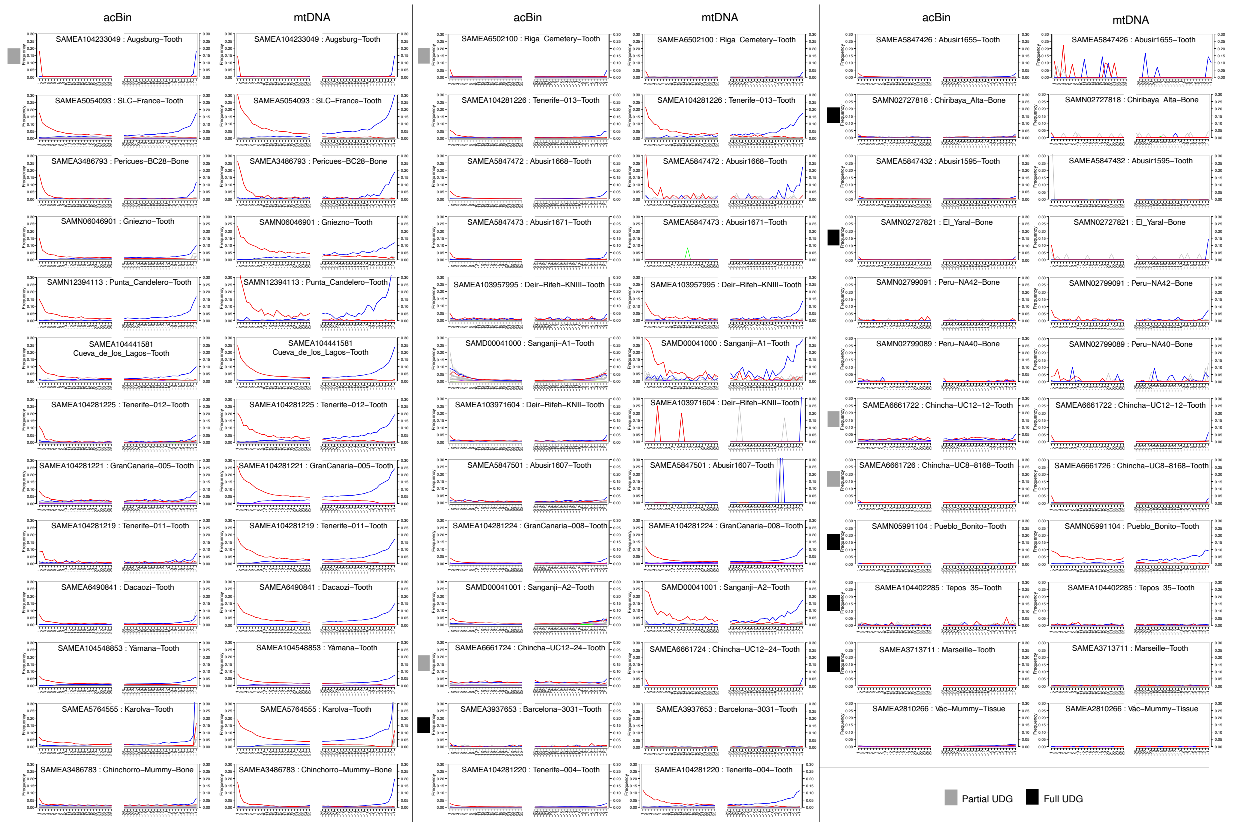


**Figure S4. MapDamage profiles depicting misincorporation levels for the first and last 25 bases of *C. tetani* and human mtDNA fragments from 38 ancient DNA samples.** G→A misincorporations (blue); C→T misincorporations (red). The top of each column is labeled by DNA type (columns 1,3,5 – acBins; columns 2,4,6 – human mtDNA) and each plot has been labeled according to its sample name and BioSample ID. Many ancient samples show a characteristic pattern of increased C→T misincorporations at the 5' end and complementary G→A mutations at the 3' end of sequence fragments. Samples are ordered based on first position C→T misincorporation rate.

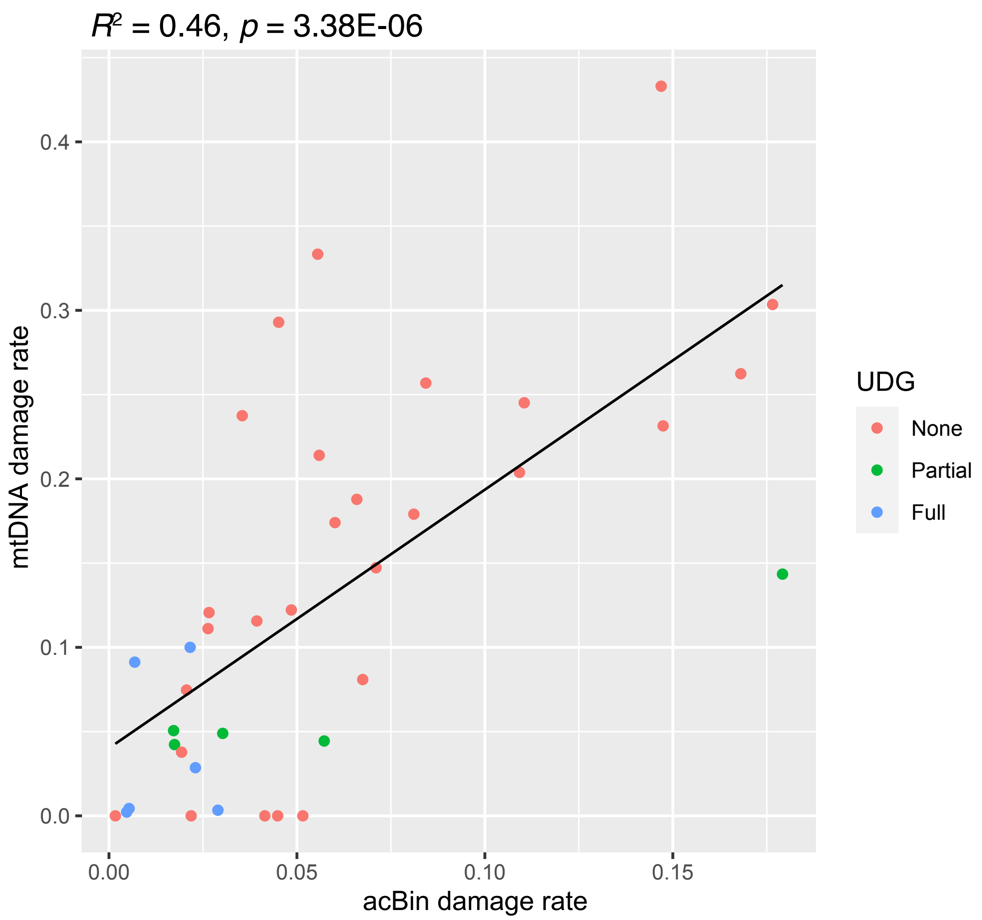


**Figure S5. Correlation between damage rates of acBins and corresponding human mtDNA from the same sample.** The acBins are also colored based on the UDG treatment used in the original aDNA samples. The damage rates are the 5' first position C→T misincorporation rate determined using mapDamage. Raw data is available in Table S2 and Table S7.

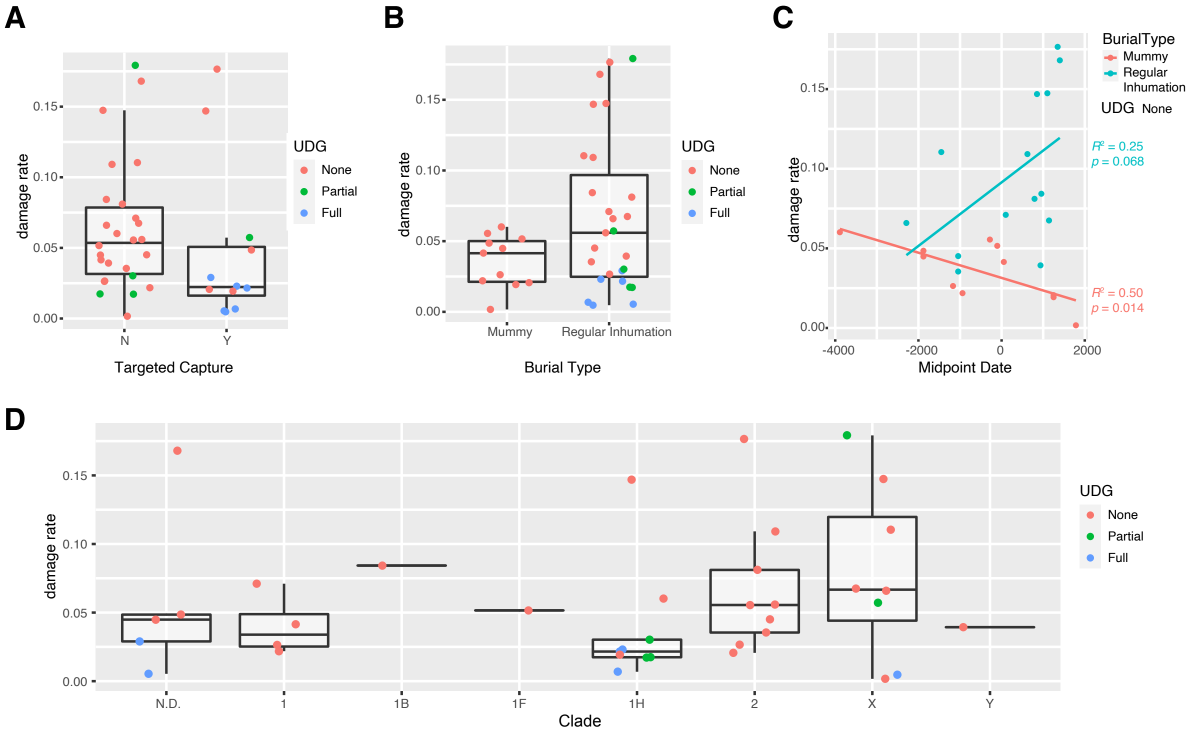


**Figure S6. Exploration of acBin damage levels versus various sample metadata.** Damage levels are plotted as a function of: A) phylogenetic clade; B) whether capture methods used; C) burial method; and D) midpoint date of archaeological sample. For plots in A, B, and D, samples are also colored according to their UDG status, as it is a main factor that influences the damage level. For the plot in C, samples with full or partial UDG treatment were removed as this is known to affect damage rates. In D, The “N.D.” group includes low coverage acBins that could not be phylogenetically placed.

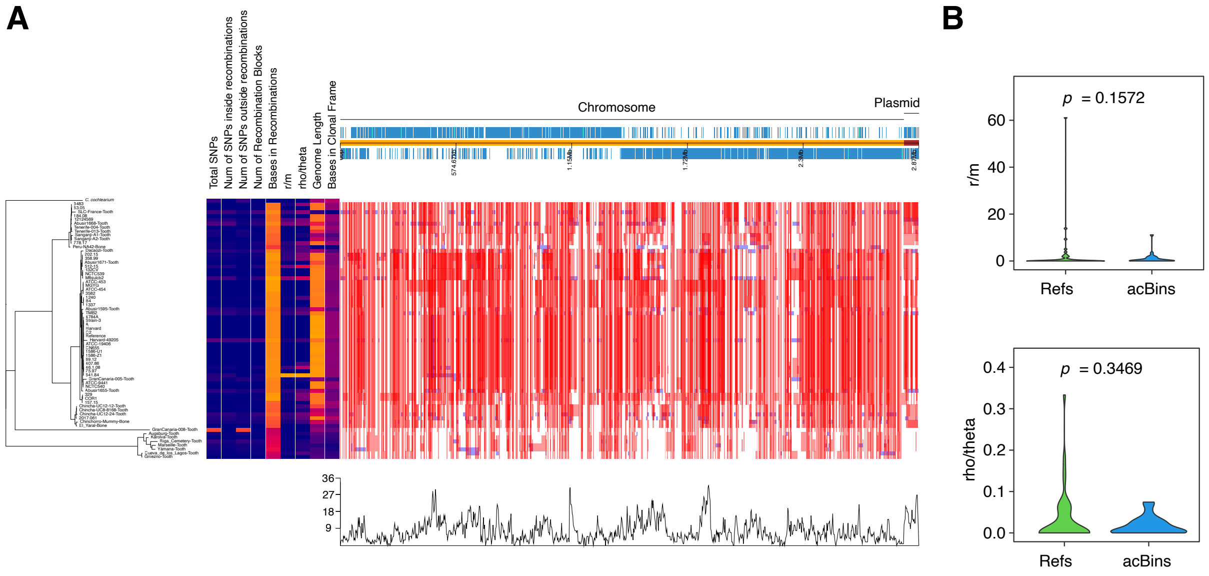


**Figure S7. Analysis of recombination using Gubbins and Phandango.** (A) The multiple alignment of acBin-derived contigs and modern *C. tetani* strains was analyzed using Gubbins and visualized using Phandango. The genome dendrogram is shown on the left and the recombination profiles are shown on the right relative to the chromosome and plasmid. The chromosome and plasmid are concatenated together to simplify the visualization. (B) Comparison of estimated recombination levels between acBins and modern strains reveals no significant differences. The r/m ratio is the ratio of the probability that a site was altered by recombination (r) and mutation (m). The rho/theta ratio measures the rate of recombination (rho) relative to the rate of mutation (theta). Genomes with gap content of 90% or greater were excluded.

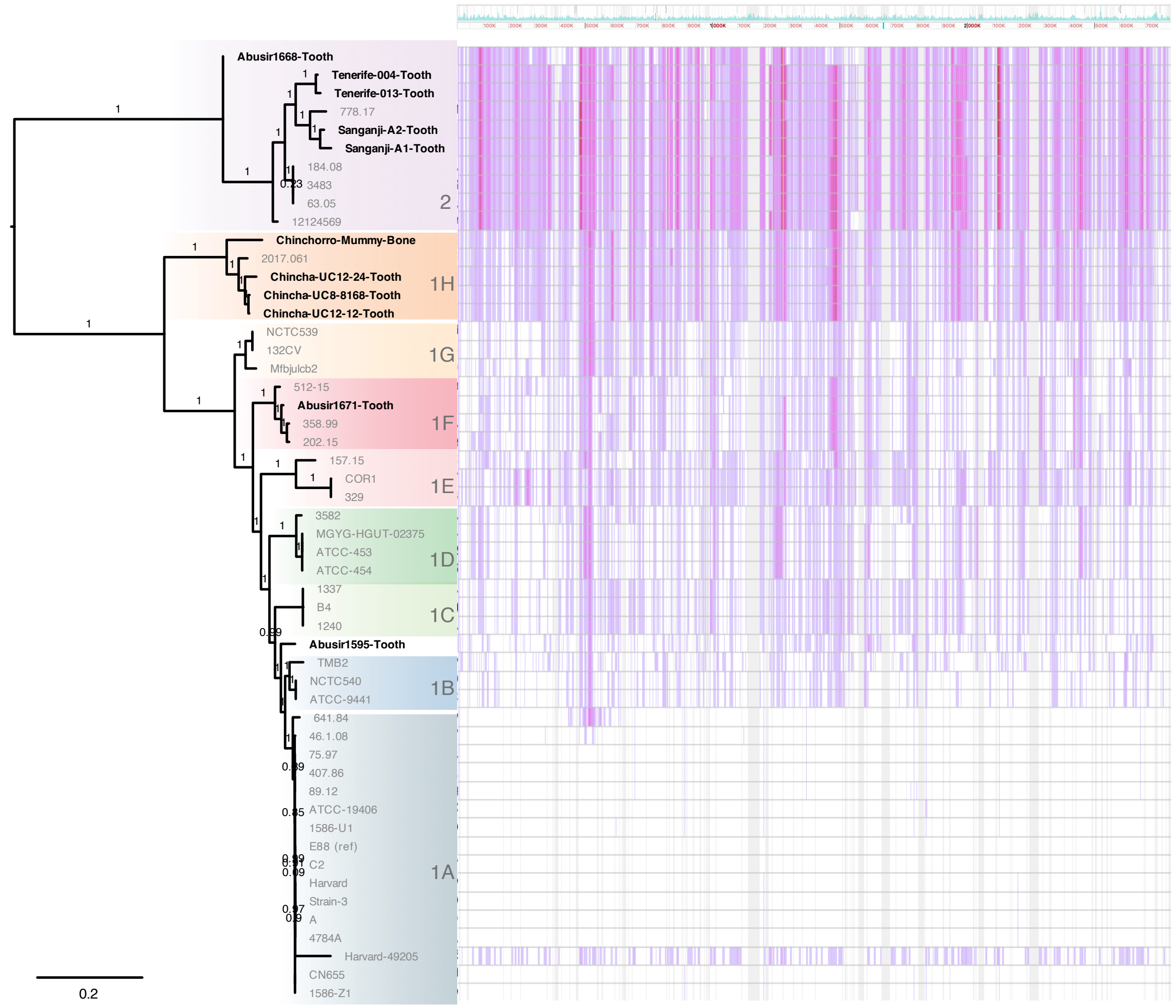


**Figure S8. SNP-based phylogenetic tree of a subset of high-coverage acBins from *C. tetani* lineages 1 and 2.** The phylogenetic tree, labeled by *C. tetani* lineage, is shown on the left with acBins shown in bold text. A visualization of the corresponding alignment using the Gingr tool is shown on the right relative to the E88 chromosome. Clade support (aLRT) values are indicated above the nodes in the tree.

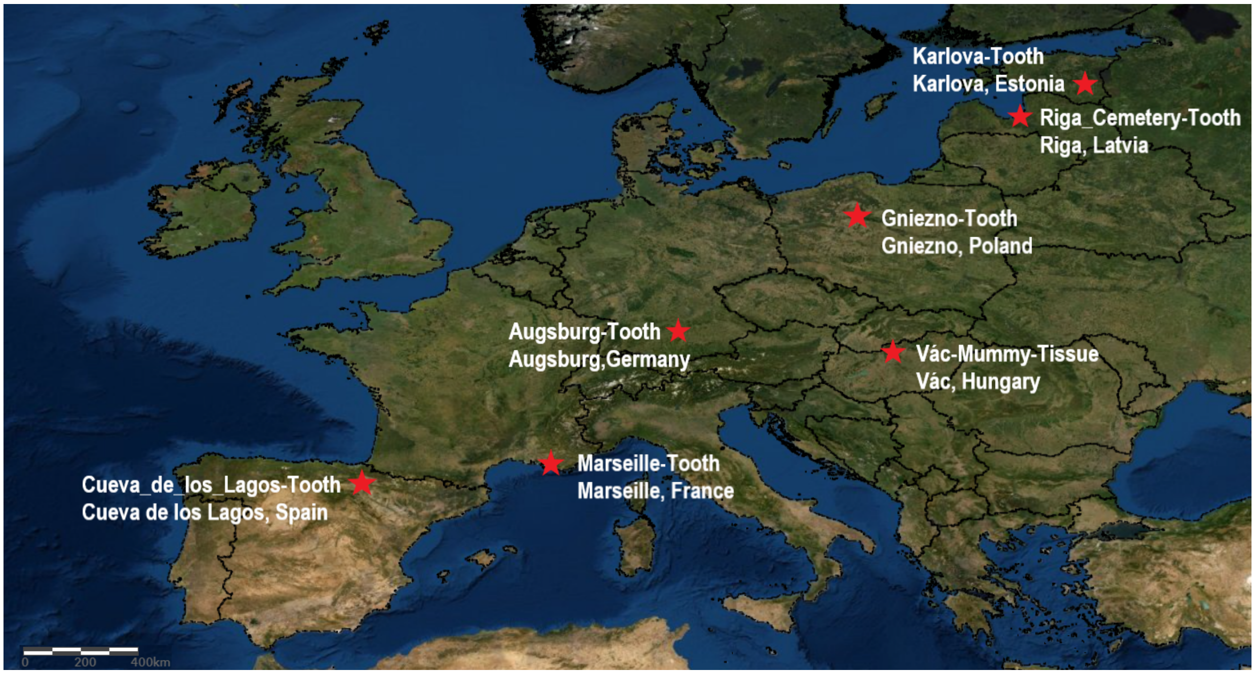


**Figure S9. Geographic location of 7/8 archaeological samples associated with *Clostridium* sp. X.** This map was made by the authors using ArcGIS online, with the World Imagery (WGS84) basemap credited to: Esri, Maxar, Earthstar Geographics, and the GIS User Community; as well as the Europe_Countries_2017 layer, credited to: Esri, and Michael Bauer Research GmbH.

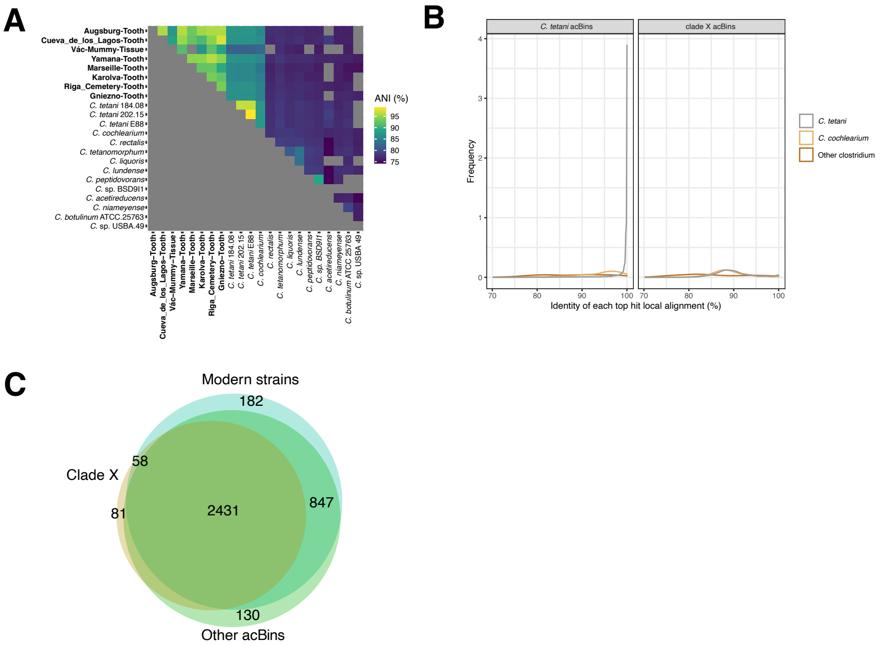


**Figure S10. Sequence similarity levels between clade X acBins and reference *Clostridium* genomes.** (A) Pairwise genome-wide average nucleotide identities (ANIs) involving acBins, representative C. tetani genomes, and other genomes from *Clostridium* species. (B) Density plot of sequence identities in local alignments between acBins and reference genomes (*C. tetani*, *C. cochlearium*, other *Clostridium* species), subdivided by clade X acBins and non-clade X acBins. (C) Venn diagram indicating overlap of predicted orthogroups between acBins, clade X, and modern *C. tetani* genomes.

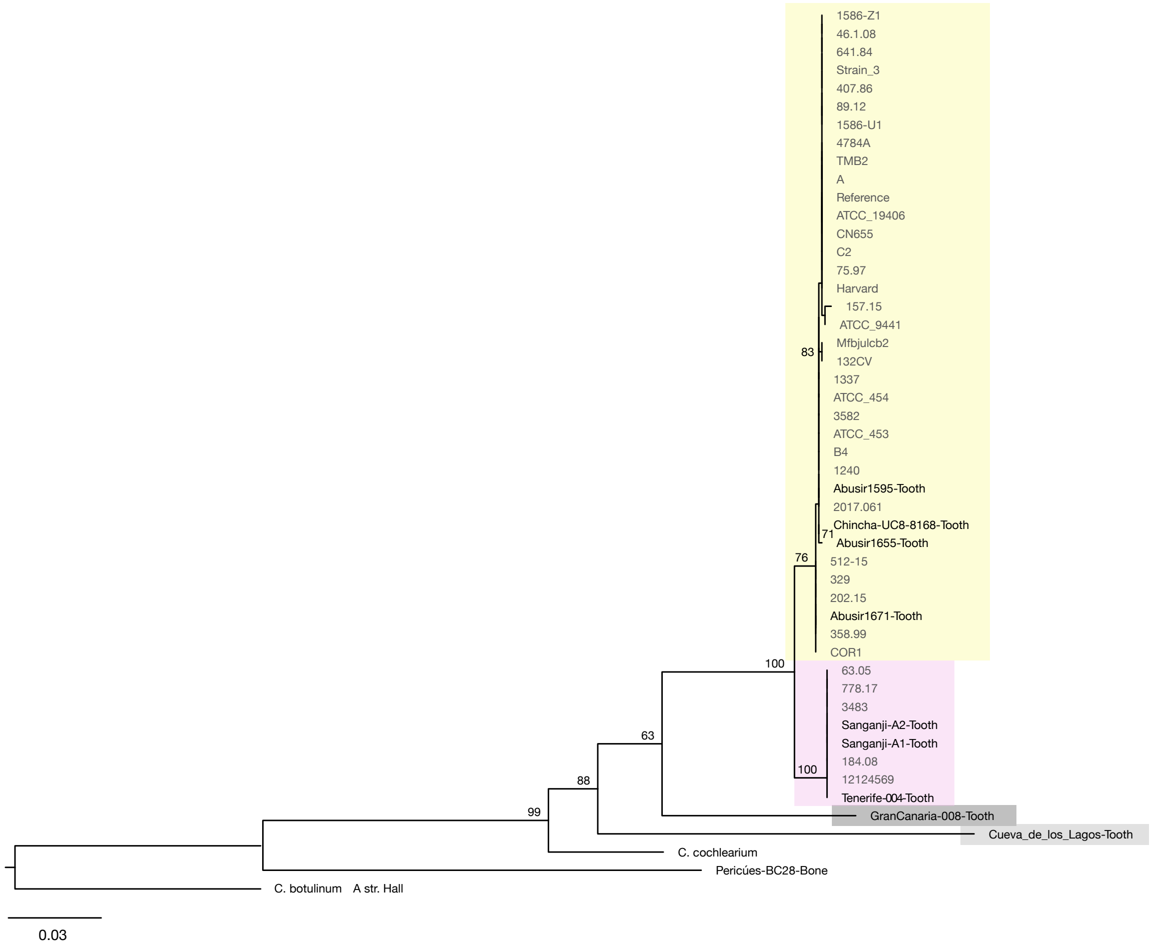


**Figure S11. Phylogenetic tree of *rpsL* coding sequences.** Trees are based on a blastn search with *Clostridium tetani* E88 sequences. The phylogeny is based on a multiple alignment of *rpsL* (AE015927.1:c2752816-2752442) sequences identifed from aDNA *C. tetani* contigs and modern *C. tetani* strains. Genome IDs of modern *C. tetani* strains are listed in Chapeton-Montes *et al*. (2019). Sequences with 80% or greater coverage of the *C. tetani* E88 query sequences were aligned with MUSCLE v3.8.31, and RAxML (v8.2.4) trees using the GTR+GAMMA model were created. Bootstrap values (based on 100 runs) are displayed for major clades. The tree is highlighted as follows: GranCanaria-Tooth008 (dark grey) and clades 1 (yellow), 2 (purple), and X (light grey).

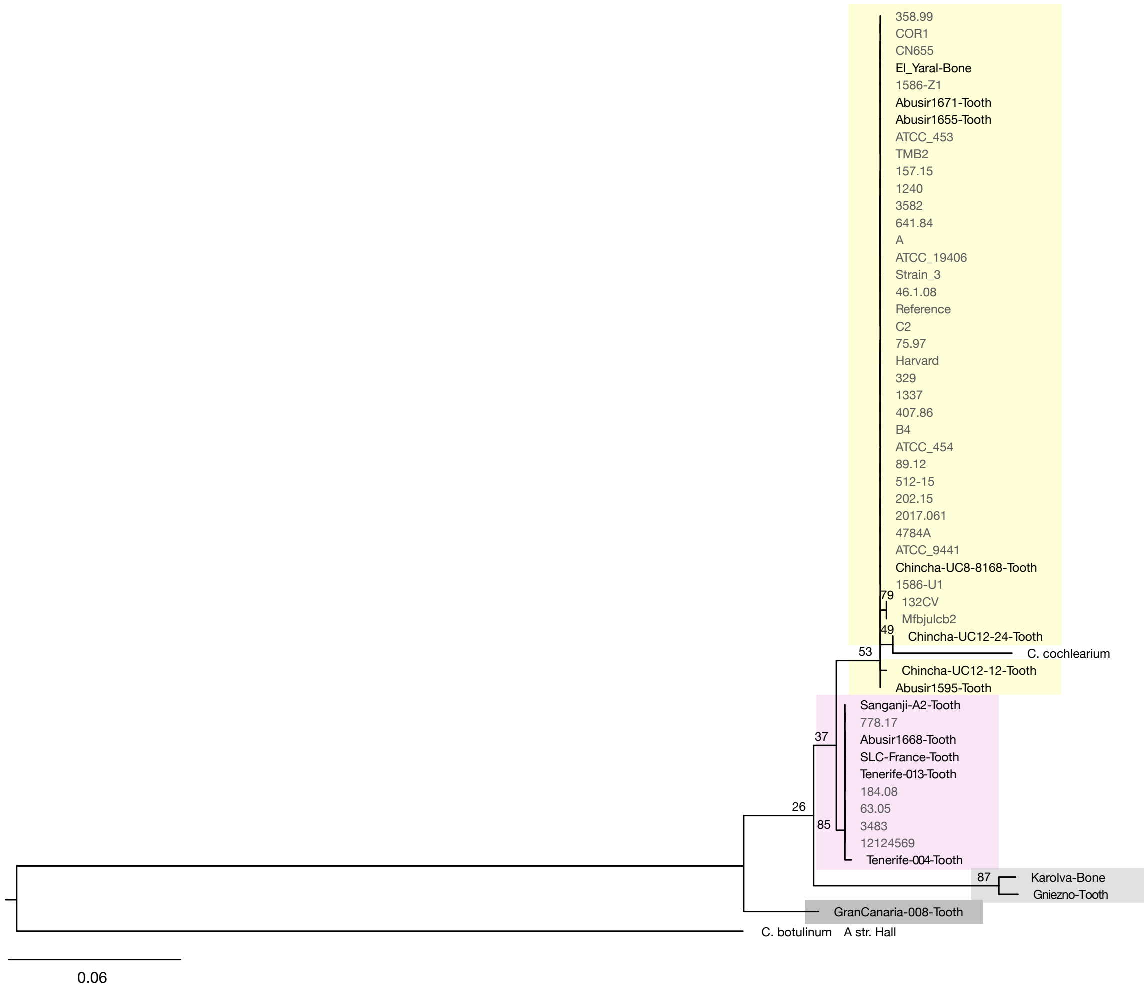


**Figure S12. Phylogenetic trees of *rpsG* coding sequences.** Trees are based on a blastn search with *Clostridium tetani* E88 sequences. The phylogeny is based on a multiple alignment of *rpsG* (AE015927.1:c2752268-2751801) sequences identified from aDNA *C. tetani* contigs and modern *C. tetani* strains. Genome IDs of modern *C. tetani* strains are listed in Chapeton-Montes *et al*. (2019). Sequences with 80% or greater coverage of the *C. tetani* E88 query sequences were aligned with MUSCLE v3.8.31, and RAxML (v8.2.4) trees using the GTR+GAMMA model were created. Bootstrap values (based on 100 runs) are displayed for major clades. The tree is highlighted as follows: GranCanaria-Tooth008 (dark grey) and clades 1 (yellow), 2 (purple), and X (light grey).

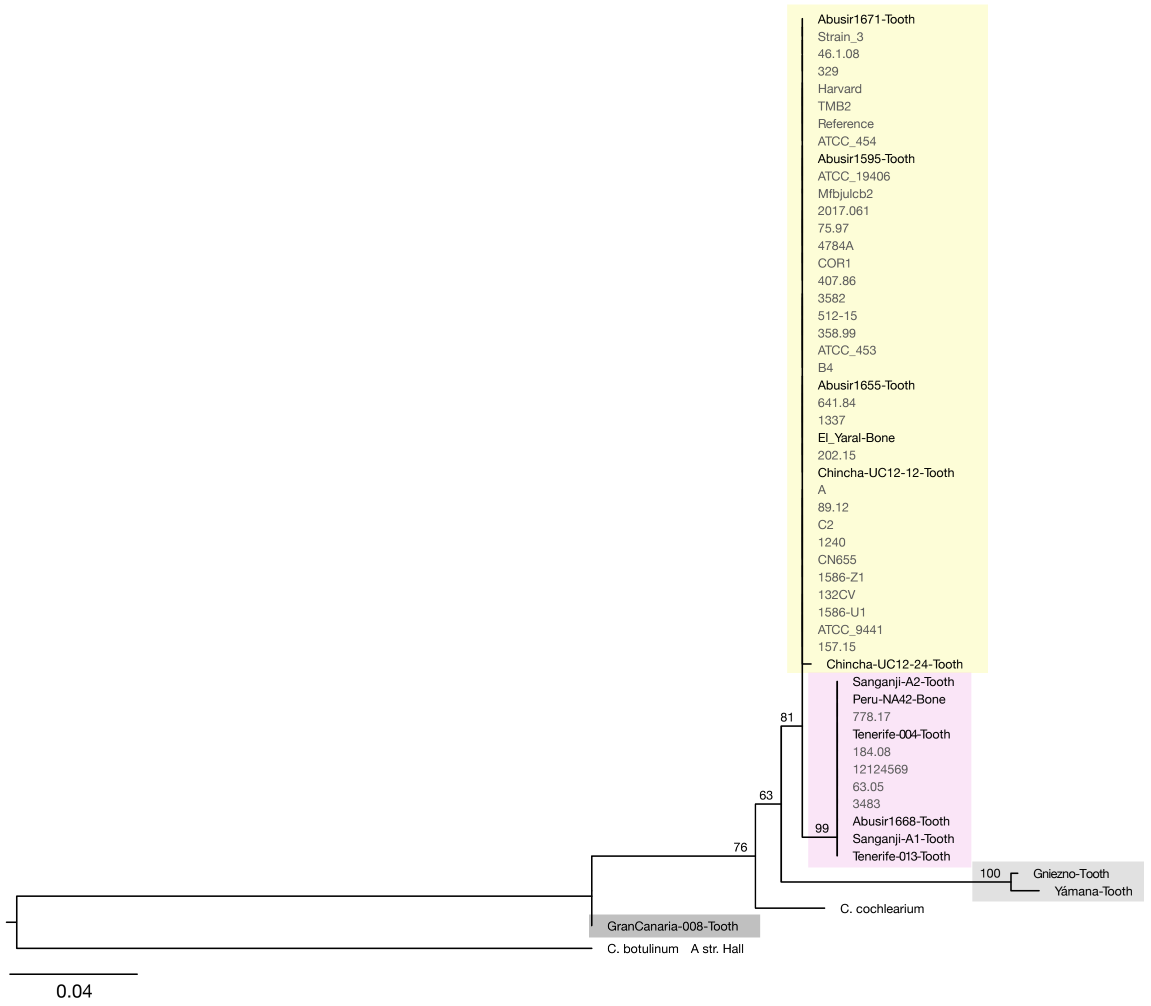


**Figure S13. Phylogenetic trees of *recA* coding sequences.** Trees are based on a blastn search with *Clostridium tetani* E88 sequences. The phylogeny is based on a multiple alignment of *recA* (AE015927.1:1383544-1384548) sequences identified from aDNA *C. tetani* contigs and modern *C. tetani* strains. Genome IDs of modern *C. tetani* strains are listed in Chapeton-Montes *et al*. (2019). Sequences with 80% or greater coverage of the *C. tetani* E88 query sequences were aligned with MUSCLE v3.8.31, and RAxML (v8.2.4) trees using the GTR+GAMMA model were created. Bootstrap values (based on 100 runs) are displayed for major clades. The tree is highlighted as follows: GranCanaria-Tooth008 (dark grey) and clades 1 (yellow), 2 (purple), and X (light grey).

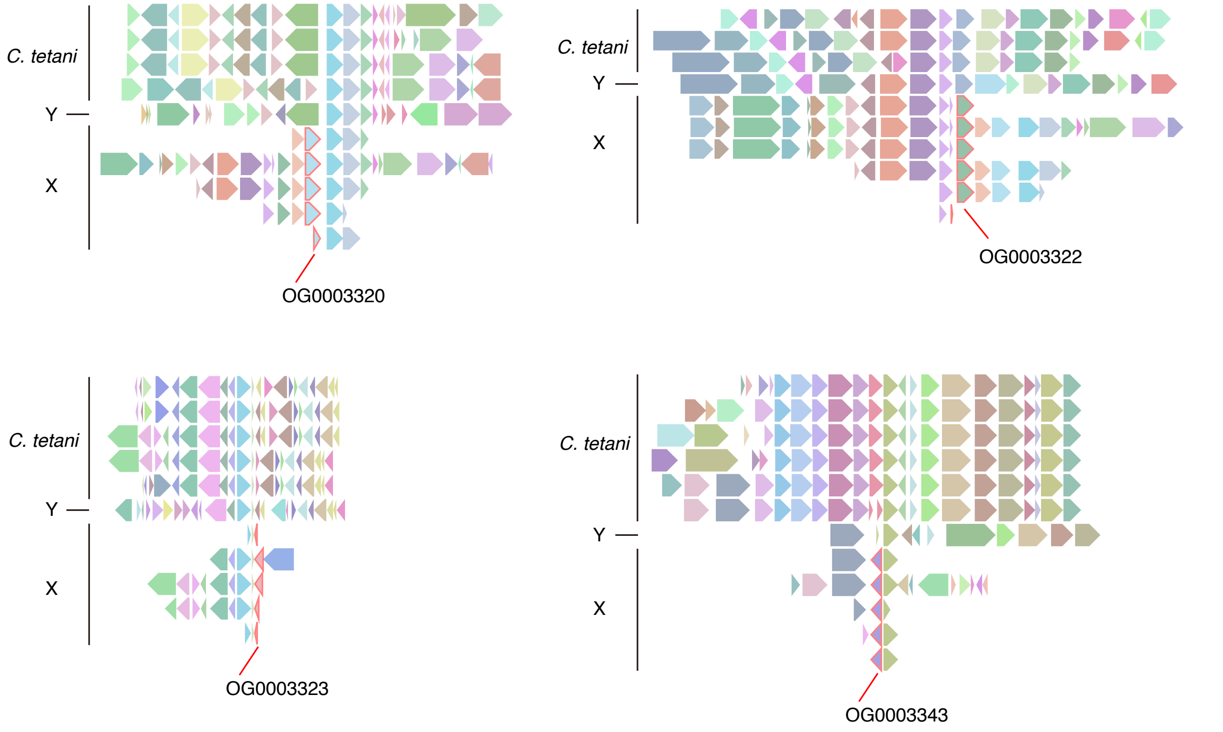


Closest BLAST matches:

OG0003320 - MYG1 family protein in *Lutibacter* sp. B2 (MBF8984673.1), 79.8 % identity
OG0003322 - EFR1 family ferrodoxin in *Desulforamulus reducens* (WP_011878543.1), 79.8 % identity
OG0003323 - SWIM zinc finger family protein in *Anaerophilus nitritogenes* (WP_129596215.1), 83.6 %ID
OG0003343 - No detected homologs

**Figure S14. Unique gene neighborhood structures in clade X and Y acBins compared to modern reference *C. tetani* strains.** Orthology-based clustering of all protein sequences from our dataset identified fourteen orthogroups (genes) unique to four or more clade X members and absent among all *C. tetani* genomes. The genomic context of four of these genes (labeled by their orthogroup ID) is shown above. The highlighted genes in red are unique to clade X. The genes surrounding these X-specific genes are conserved in other *C. tetani* genomes.

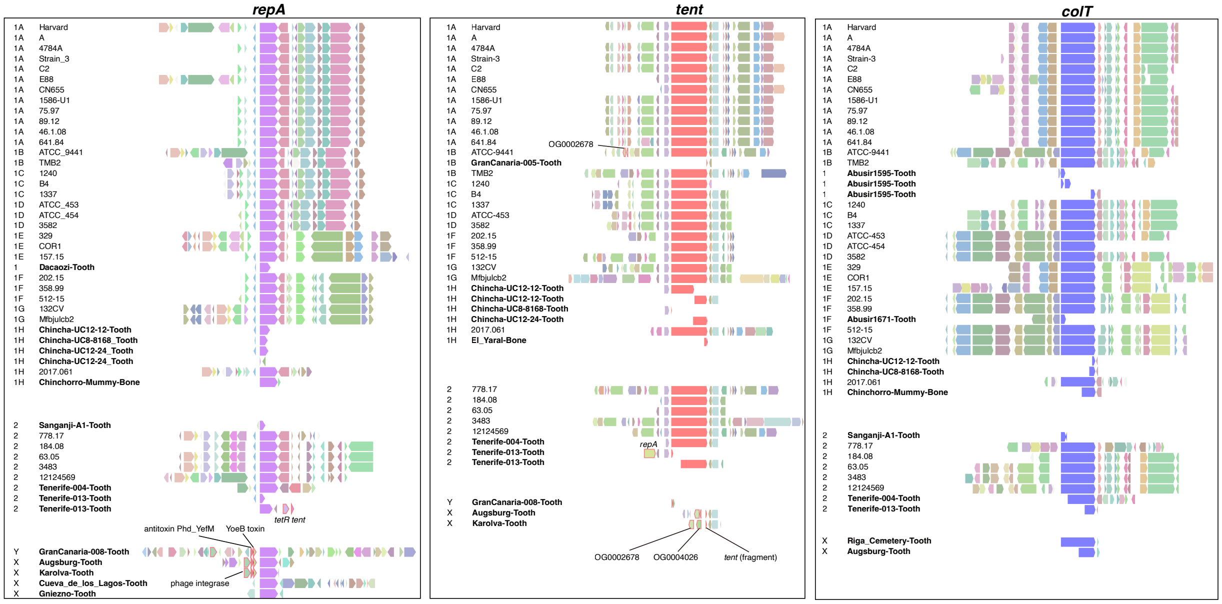


**Figure S15. Gene neighborhoods surrounding the *repA*, *colT, and tent* genes in select acBins and reference *C. tetani* genomes.** Only assembled gene clusters with one or more genes surrounding the center gene were selected for display. Genes are colored uniquely based on their orthogroup membership as determined by OrthoFinder predictions. Gene clusters from *C. tetani* lineage 1, *C. tetani* lineage 2, and *Clostridium* sp. X and Y are grouped separately. In general, gene clusters from *C. tetani* lineage 1 and 2 share similarities, and those from *Clostridium* sp. X and Y contain patterns that are unique from *C. tetani*. On the left acBins names are bolded.

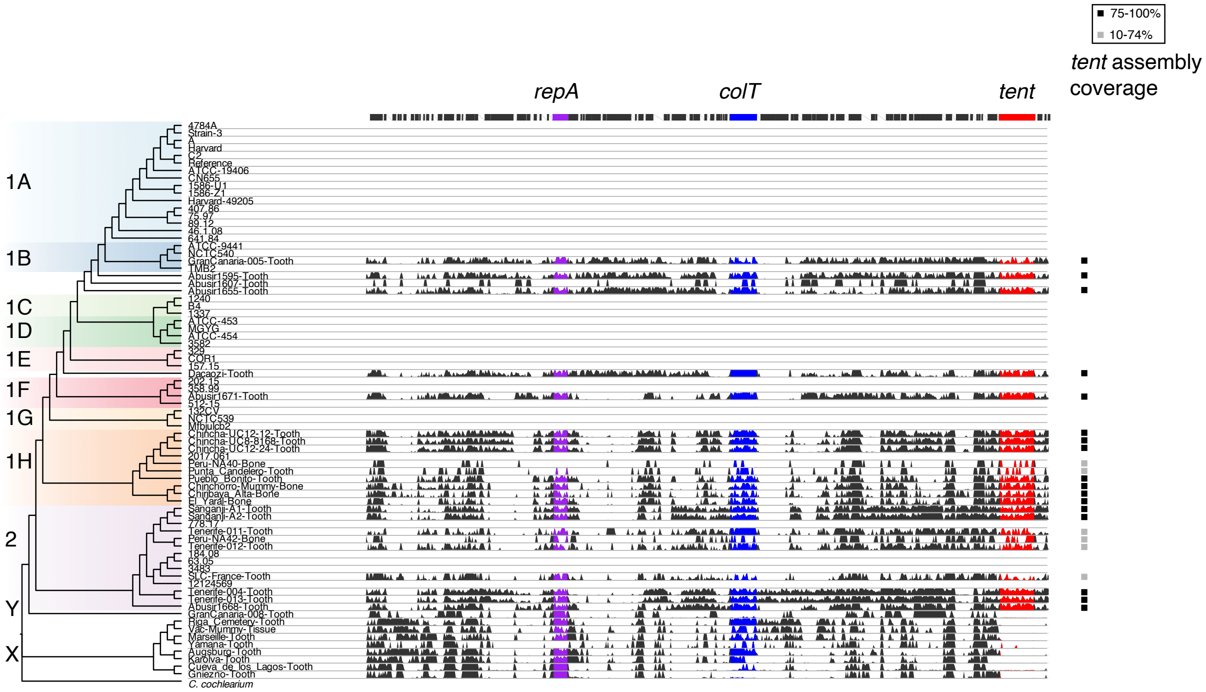


**Figure S16. Mapped read coverage to the reference *C. tetani* E88 plasmid and *tent* genes.** A) Visualization of mapped read coverage to the *C. tetani* E88 plasmid. The genes (*tent*, *colT,* and *repA*) are coloured red, blue and purple, respectively. The plot reveals a relative lack of coverage in clade X acBins across the *tent* gene. Coverage values were capped to the 80th percentile to prevent high coverage regions from obscuring other regions. Genes were plotted as black bars using RefSeq annotations. B) Normalized abundance of *tent* detected per acBin clade. The coverage (average sequencing depth) of reads mapped to the *tent* gene was normalized to the coverage of the *repA* gene, a plasmid marker. On the right of the plot, the boxes indicate the acBins for which *tent* sequences were assembled with high coverage (black) or partially (gray) using SNP profiling as described in the Methods.

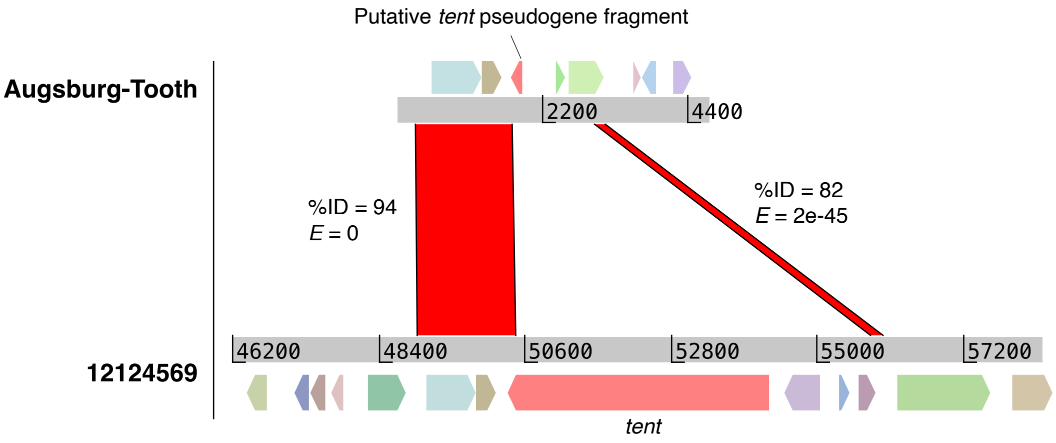


**Figure S17. Identification of a putative deletion event resulting in a possible *tent* pseudogene in clade X acBins.** Two clade X acBins possess an assembled contig containing a *tent* fragment as shown in Figure 3D. A BLASTN comparison of this contig from Augsburg-Tooth against a representative *tent* locus from strain 12124569 reveals two regions of homology that flank the *tent* gene in strain 12124569. This indicates a possible rearrangement or deletion that has resulted in the removal of the *tent* gene and some adjacent sequence in Augsburg-Tooth.

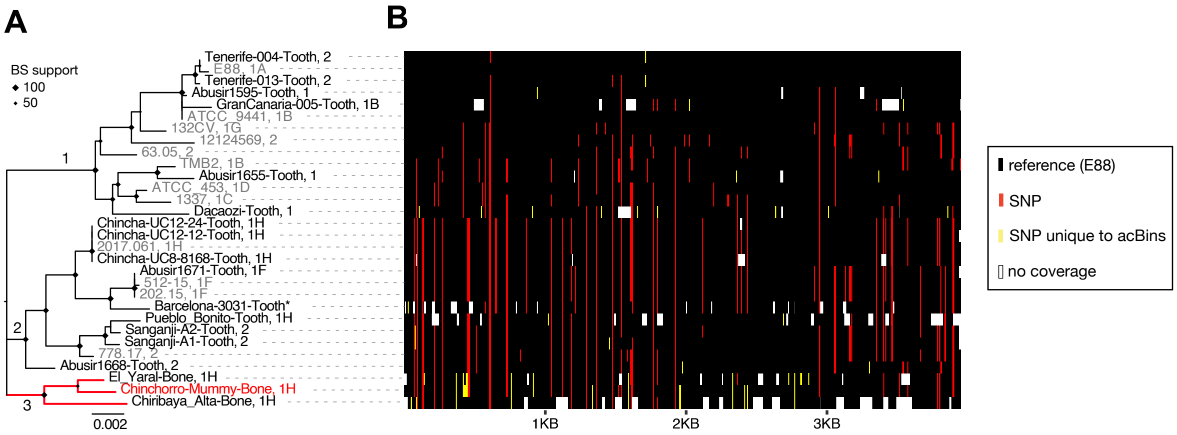


**Figure S18. Reconstructed *tent* sequences from ancient DNA samples display unique mutational profiles from modern sequences.** Maximum-likelihood phylogenetic tree of *tent* genes (left) and visualization of SNP profiles relative to the E88 reference sequence (right). The visualization on the right plots the SNP variation in each sequence, with yellow positions representing SNPs found uniquely in *tent* sequences from ancient DNA samples, red positions indicating SNPs also found in modern *tent* sequences, and black positions indicating identity to the reference E88 *tent* sequence. Gaps are colored white. The plot was generated by loading the *tent* MSA into R v4.1.0 with the Biostrings library v2.60.1, converting the MSA to a data matrix using a custom script (see Github repository) and plotted as a tile plot using the ggplot2 library v3.3.3.

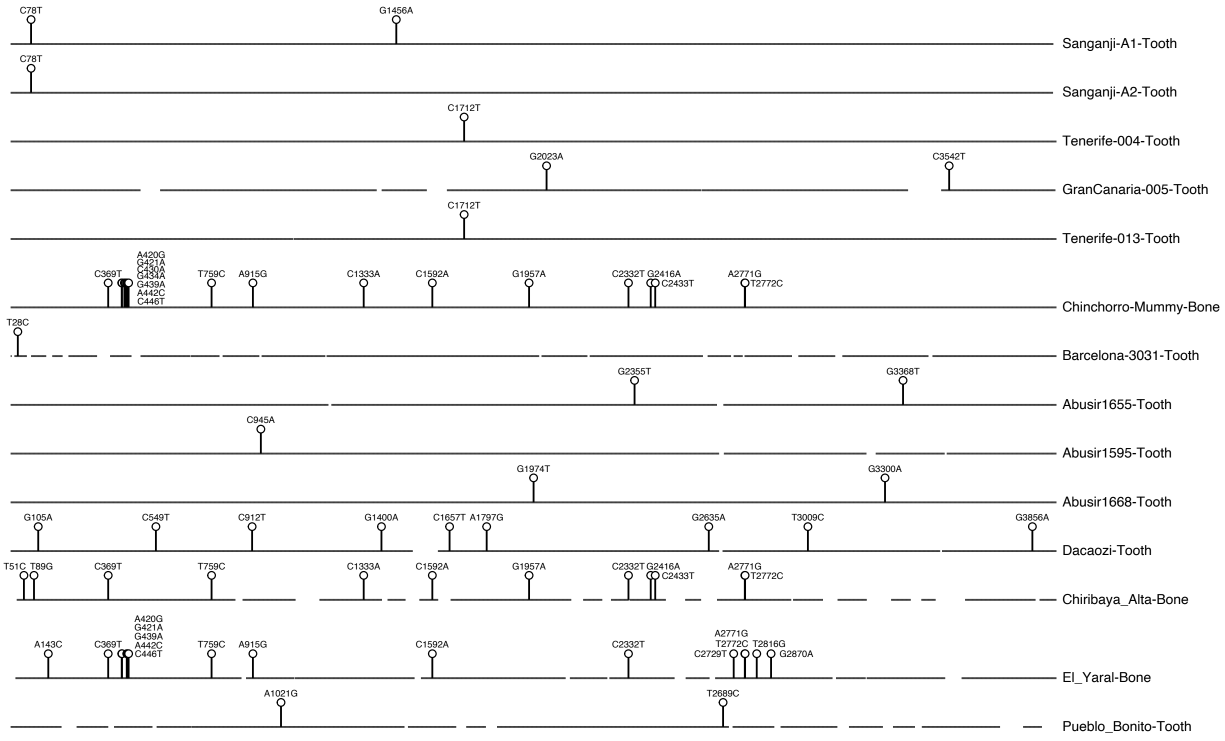


**Figure S19. Novel nucleotide substitutions in *tent* genes recovered from ancient samples that have not been identified in modern *tent* genes.** Each sequence is depicted in their 5'-3' orientation. Horizontal lines indicate sequences that are identical to the E88 reference *tent* sequence, and missing lines indicate gaps (regions with no coverage). All SNPs are listed with numbering relative to the E88 reference *tent* sequence.

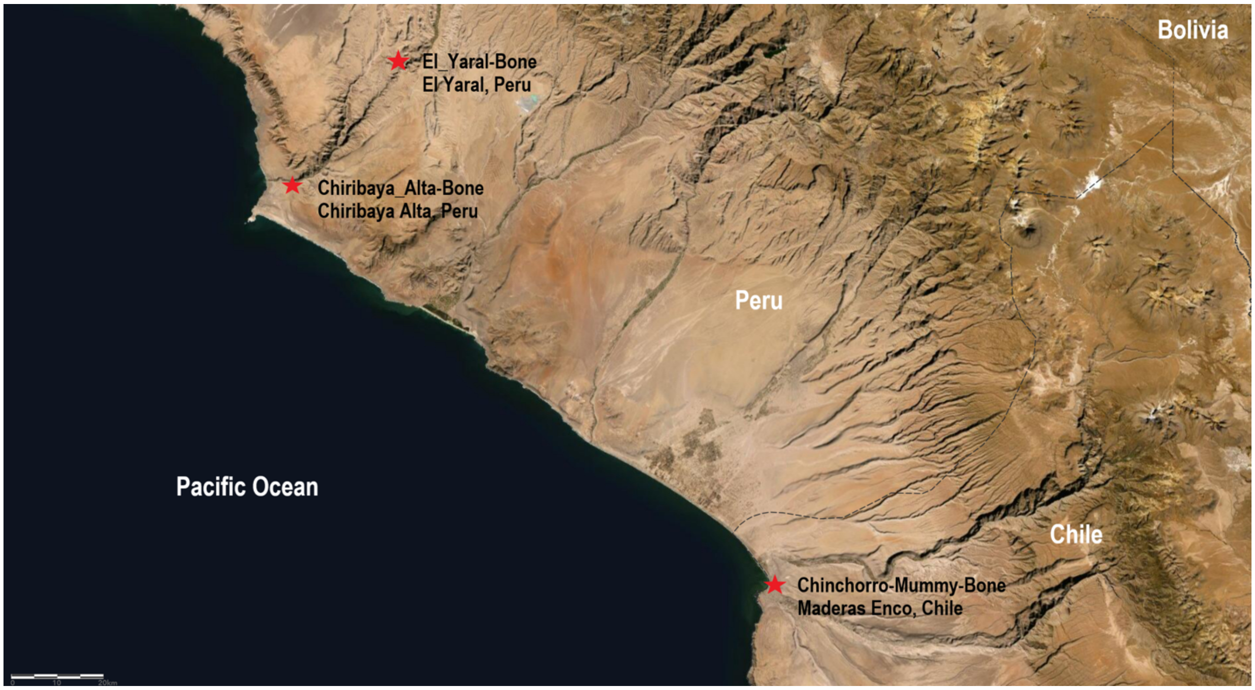


**Figure S20. Geographic location of three samples [Chinchorro-Mummy-Bone (SAMEA3486783), El_Yaral-Bone (SAMN02727821), and Chiribaya_Alta-Bone (SAMN02727818)] associated with subgroup '2' *tent* variants**. This map was made by the authors using ArcGIS online, with the World Imagery (WGS84) basemap credited to: Esri, Maxar, Earthstar Geographics, and the GIS User Community; as well as the Chile Country Boundary 2020 and Bolivia Country Boundary 2020 layers, both credited to: Esri, Michael Bauer Research GmbH 2020, Instituto Nacional de Estadísticas, UN.

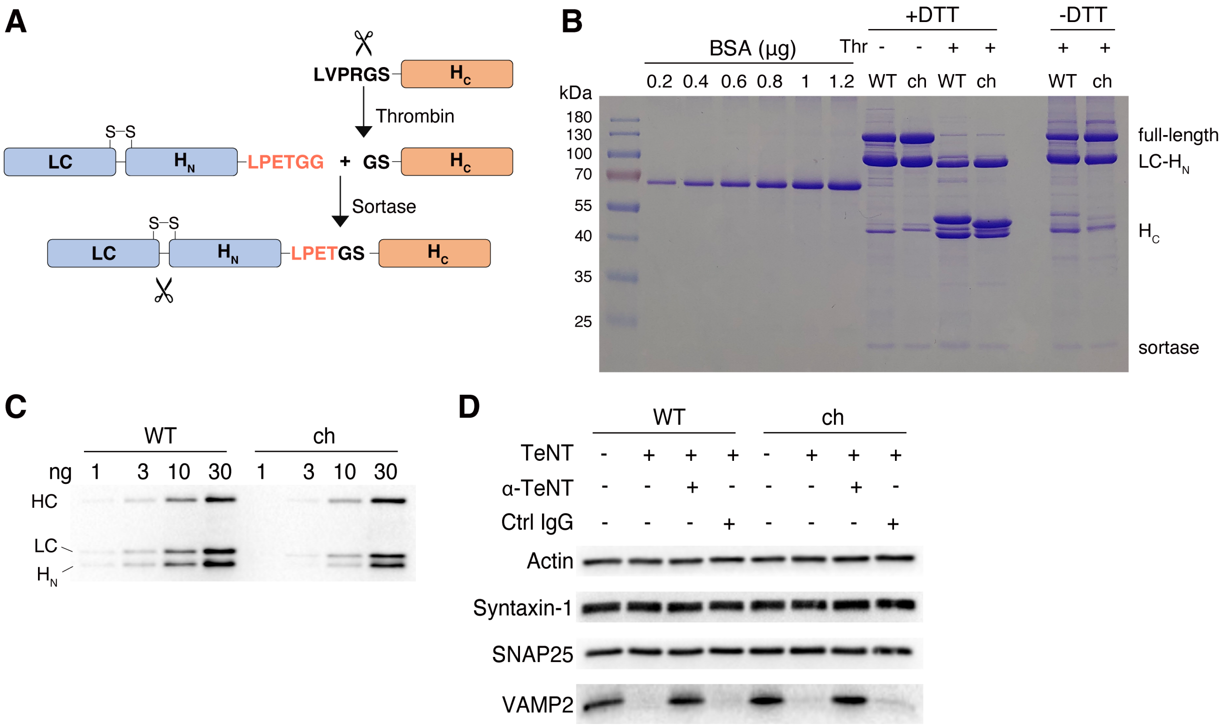


**Figure S21. Antiserum of TeNT neutralized TeNT and TeNT/Chinchorro with similar efficiency.**
(A) The schematic drawing of sortase ligation method.
(B) Sortase ligation reaction mixtures were analyzed by SDS-PAGE and Coomassie blue staining. Thr, Thrombin.
(C) Rabbit antiserum of TeNT recognized full-length activated TeNT and TeNT/Chinchorro ("ch") with similar sensitivity by using western blot analysis.
(D) Cultured rat cortical neurons were exposed to 25 pM full-length toxins together with antiserum (α-TeNT, 1:1000) or control non-immunized serum (Ctrl IgG, 1:1000) for 12 hrs. Cell lysates were analyzed by immunoblot.

**Table S1**. Metadata for SRA sequencing runs associated with high *C. tetani* abundance predicted using the SRA taxonomy analysis tool.
**Table S2**. Metadata and bioinformatic annotations for BioSamples containing ancient human DNA and C. tetani. Metadata includes SRA-derived information as well as newly added annotations based on analysis of acBins.
**Table S3**. Identified archaeal and bacterial species in *C. tetani*-containing human aDNA samples. Total counts for each biosample are derived from their respective SRA taxonomic profiles.
**Table S4**. Kaiju taxonomic classification summary at the species level of assembled samples (contig count)
**Table S5.** Statistics for assembled contigs assigned to *C. tetani* after blast-validation
**Table S6**. CheckM analysis of all 38 acBins blast-verified C. tetani contigs
**Table S7**. MapDamage analysis of acBin contig sets. Contigs were BLAST-validated as mapping to C. tetani and contigs encoding rRNAs and tRNAs were removed.
**Table S8**. PyDamage analysis of acBin contig sets. Contigs were BLAST-validated as mapping to C. tetani and contigs encoding rRNAs and tRNAs were removed.
**Table S9**. Pairwise average nucleotide identity values for a *C. tetani* bin from a human gut sample (SRR10479805) to acBins and modern genomes from related *Clostridium* species.
**Table S10**. Pairwise average nucleotide identity values for *Clostridium* sp. clade X acBins and modern genomes from related *Clostridium* species.
**Table S11**. Pairwise average nucleotide identity values for lineage Y acBin (GranCanaria-008-Tooth) and modern genomes from related *Clostridium* species.
**Table S12**. Per-sample coverage statistics and correlations for the *C. tetani* chromosome, plasmid, *repA*, *colT* gene, and *tent* gene.
**Table S13**. Comparison of assembled *tent* sequences from ancient DNA samples with modern *tent* sequences.
**Table S14**. Detected SNPs in tent sequences from acBins and modern C. tetani strains relative to tent/E88 reference.
**Table S15**. *tent* variants detected in ancient samples using the Octopus variant caller. See VCF file format for header information.
**Table S16**. Analysis of SNPs present in tent sequences from Chinchorro_Mummy_bone, El_Yaral_Bone, and Chibaya_Alta-Bone
**Table S17**. FASTQ processing statistics for ancient DNA samples with high predicted *C. tetani* content.
**Table S18**. Genomes of *C. tetani* strains downloaded from the NCBI database and associated metadata.
